# Synonymous mutations reveal genome-wide driver mutation rates in healthy tissues

**DOI:** 10.1101/2020.10.08.331405

**Authors:** Gladys Poon, Caroline J. Watson, Daniel S. Fisher, Jamie R. Blundell

## Abstract

Genetic alterations that drive clonal expansions in ostensibly healthy tissues have implications for cancer risk. However, the total rate at which clonal expansions occur in healthy tissues remains unknown. Synonymous passenger mutations that hitchhike to high variant allele frequency due to a linked driver mutation can be used to estimate the total rate of positive selection across the genome. Because these synonymous hitchhikers are influenced by *all* mutations under selection, regardless of type or location, they can be used to estimate how many driver mutations are missed by narrow gene-focused sequencing panels. Here we analyse the variant allele frequency spectrum of synonymous passenger mutations to estimate the total rate at which mutations driving clonal expansions occur in healthy tissues. By applying our framework to data from physiologically healthy blood, we find that a large fraction of mutations driving clonal expansions occur outside of canonical cancer driver genes. In contrast, analysis of data from healthy oesophagus reveals little evidence for many driver mutations outside of those in *NOTCH1* and *TP53*. Our framework, which generalizes to other tissues, sheds light on the fraction of drivers mutations that remain undiscovered and has implications for cancer risk prediction.

## Introduction

Next-generation sequencing of healthy tissues has revealed that large numbers of non-synonymous mutations are under positive selection and cause clonal expansions ^1–11^. These expanded ‘driver’ mutations, which often occur in cancer-associated genes, have an increased chance of acquiring further pathogenic mutations and thus have implications for cancer risk ^1,2,10–13^. Clonal expansions are typically identified via high-depth sequencing of a ‘panel’ of cancer-associated genes ^4,5,7,14,15^. However, because a large fraction of the genome lies beyond the target regions of these sequencing panels and the panels are often designed to best detect single nucleotide variants and indels, mutations in non-coding regions and more complex mutations including mosaic chromosomal alterations (mCAs) ^10,11^ and structural variants (SVs) are potentially missed. Moreover, the fact that some cancers are found with no known driver mutations suggests there might be a large number of drivers which are individually rare but collectively common ^6^. This raises the question: how many mutations driving clonal expansions are missed by gene-focused sequencing panels?

Positive selection for a gene or variant is commonly identified using recurrence (e.g. over representation of mutation in a particular gene) ^16^, elevated ratios of nonsynonymous to synonymous mutations (dN/dS) ^4,6^ and analysis of the distribution of variant allele frequencies (VAF) ^17,18^. However none of these approaches naturally extend to quantifying the *total* rate of positive selection. Recurrence misses mutations that are rare ^16^ and confounds the effects of selection with mutation (e.g. recurrence can be high at mutational hotspots) ^19^. Approaches based on dN/dS are restricted to genes ^4,6^. Methods based on the variant allele frequency spectrum can disentangle selection from mutation and are not restricted to genes but these also require large numbers of individuals to share the same mutation to achieve accurate estimates ^17^. Therefore estimating the total burden of positive selection outside of genes using existing methods is challenging.

Here we develop a population genetic framework that can estimate the *total* genome-wide rate of positive selection by considering the distribution of VAFs of synonymous variants. We show that most synonymous variants reach high VAF due to genetic hitchhiking: they are ‘passenger’ mutations that co-occur with a positively selected ‘driver’ mutation which itself might be undetected. The number of these high VAF synonymous variants thus provides information about the genome-wide rate of ‘driver’ mutation. Specifically, each ‘driver’ mutation will generate a ‘comet tail’ of synonymous passenger mutations as it clonally expands ^20^. Once these driver mutations reach high VAF, the VAF distribution of their synonymous passengers declines with the inverse square of the VAF, often referred to as a ‘LuriaDelbrück’ distribution ^21,22^. This characteristic distribution occurs whenever selectively neutral mutations are acquired in an exponentially growing population ^20,23–27^. The number of passenger mutations generated in a single ‘driver’ event is typically small. However, by aggregating measurements of all synonymous variants across many samples of the same tissue, a statistical distribution of synonymous passenger mutations emerges that can be used to estimate the total rate at which ‘driver’ mutations are acquired in that tissue. Since synonymous variants can hitchhike with *any* positively selected mutation, regardless of type or location in the genome, this approach provides an estimate for the *total* rate of positive selection in the genome, including non-coding mutations and more complex genetic and epigenetic alterations.

Applying our framework to data from healthy blood ^14,15^, we find evidence of a clear transition whereby low VAF synonymous variants are dominated by genetic drift while high VAF synonymous variants are predominantly caused by genetic hitchhiking. By using the high VAF synonymous passengers to quantify the total rate of positive selection across the genome, we estimate that ∼ 90% of clonal expansions leading to high VAF synonymous passengers in blood are driven by mutations outside of canonical cancer-associated genes. Applying the same framework to data from healthy oesophagus ^5^, we estimate that ∼ 60% of clonal expansions in healthy oesophagus are caused by mutations occurring in just two driver genes (*NOTCH1* and *TP53*). Our method therefore suggests that there could be many non-coding, structural and copy number ‘driver’ mutations still to be discovered in healthy blood and relatively few in healthy oesophagus.

## Results

### Inferring rate of drivers from synonymous variants

To understand how the rate and selective strength of ‘driver’ mutations shape the VAF distribution of synonymous variants, we first consider a stochastic model of stem cell dynamics (Figure 1). In the initial ‘development’ phase, stem cells grow exponentially from a single cell until reaching a fixed population size, *N*. Then, in the subsequent ‘homeostasis’ phase, stem cell numbers remain at *N* by virtue of stochastically self-renewing and terminally differentiating at the same rate 1*/τ* where *τ* is the time in years between symmetric stem cell divisions ^17^ (Figure 1A). Mutations entering the stem cell population during homeostasis are either synonymous, which are acquired at total rate *µ*_*n*_, or ‘drivers’, which are acquired at total rate *µ*_*b*_ (both per cell per year). The rate of acquiring ‘driver’ mutations is low enough that competition between two or more large ‘driver’ clones within the same individual (‘clonal interference’) is unlikely. Synonymous mutations, being neutral, do not alter the balance between self-renewal and differentiation (Figure 1A), whereas ‘driver’ mutations increase the rate of self-renewal relative to differentiation by a magnitude *s* per year, (termed the ‘fitness effect’), enabling them to exponentially expand (Figure 1B). Fitness effects of ‘driver’ mutations are drawn from a distribution (DFE) to reflect the range of functional consequences of different mutations. Synonymous mutations co-occurring in the same cell as a ‘driver’ are termed ‘passengers’ and also expand exponentially due to genetic linkage with the ‘driver’ mutation. The VAF distribution of synonymous variants that results from this process has two defining features.

**Fig. 1.**
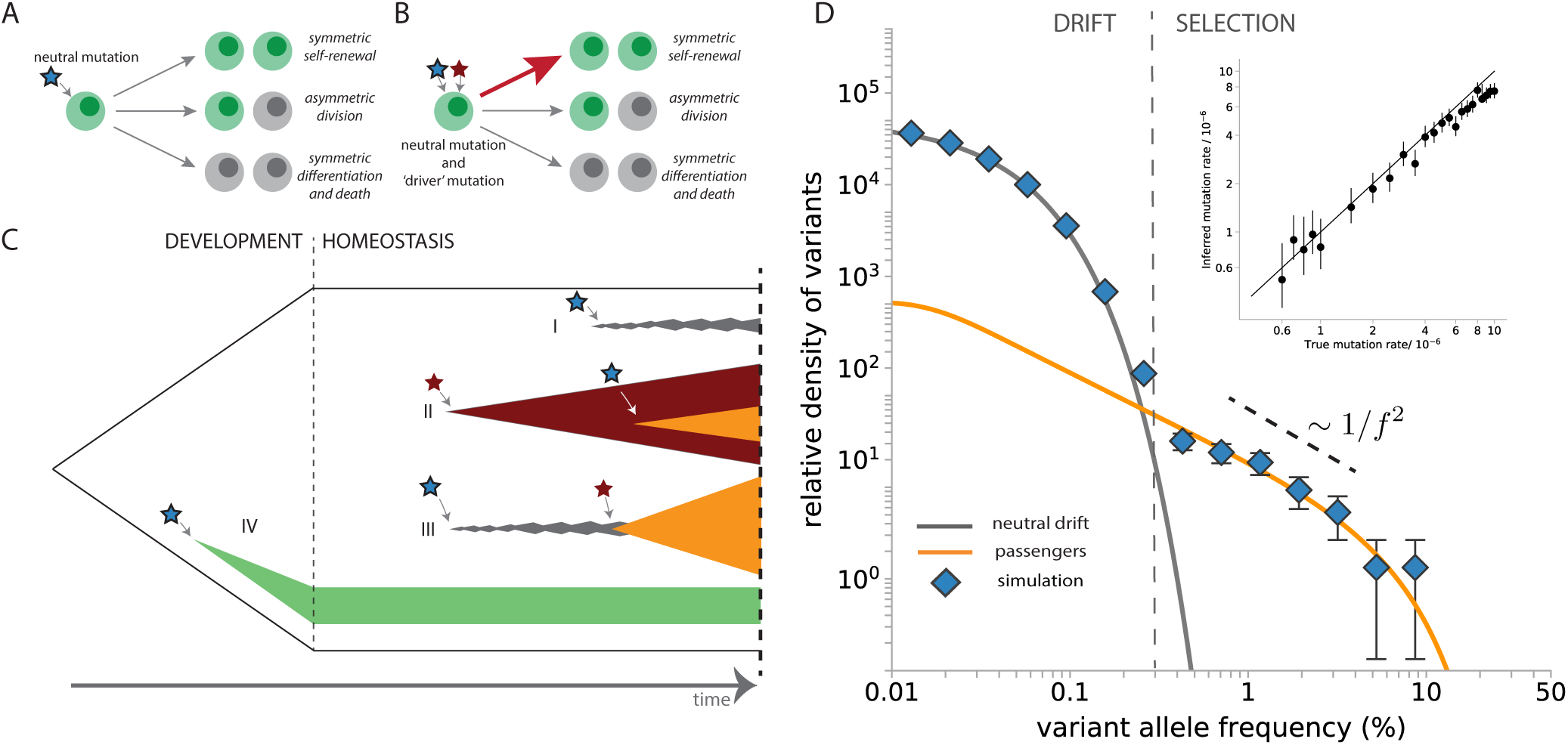
A model of genetic hitchhiking. (**A**) In a stochastic branching model of stem cell dynamics, self-renewal and differentiation rates remain balanced in cells harbouring neutral synonymous mutations. (**B**) A synonymous mutation (blue star) which co-occurs with a ‘driver’ mutation (maroon star) is termed a ‘passenger’. The ‘driver’ causes a skew in cell fates towards self-renewal which also impacts the passenger due to genetic linkage. (**C**) A schematic depicting the four different classes of synonymous mutations that are considered here: (I) a synonymous mutation arising in adulthood not linked to any driver mutation performs neutral drift and remains at low VAF. (II) an expanding clone with a ‘driver’ mutation subsequently acquires a synonymous passenger mutation which hitchhikes to high VAF. (III) a neutral clone with a synonymous mutation subsequently acquires a ‘driver’ mutation and hitchhikes to high VAF. (IV) synonymous mutations that occur early during development can reach high VAF. (**D**) VAF spectra of neutral mutations (blue data points) from stochastic simulations are overlaid with theoretical predictions for drift (‘neutral drift’, grey line) and hitchhiking (‘passengers’, orange line). The amplitude of the high VAF synonymous variants accurately recovers the true rate of driver mutations (Inset). The main figure shows an instance of simulation (Supplementary note 2) with ‘driver’ mutation rate *µ*_*b*_ = 3 × 10^−6^ per year at age 70 for the case where for *τ* = 1 year.

#### Synonymous variants at low VAF are drift-dominated

Synonymous variants that are detected at low VAF are most likely to have arisen during the homeostasis phase and to exist in clones that have not acquired a ‘driver’ mutation (Supplementary note 1A). These neutral clones are therefore subject to the forces of genetic drift alone (Figure 1C, case I) and remain concentrated at low VAFs because they do not have a fitness advantage (*s* = 0). The VAF distribution which results from these synonynous variants being generated at a constant rate and subsequently drifting scales as 1*/f* at low VAF and then falls away exponentially at VAF *> ϕ* = *t/*2*Nτ* which is proportional to age, *t* ^17,18,28^, (Supplementary note 1A). Because age, *t*, is a known quantity, the frequency at which the exponential fall-off happens, *ϕ*, provides an important independent check on the value of *Nτ*. We validated this prediction with stochastic simulations (Supplementary note 1) in which we recorded the density of all synonymous variants as a function of VAF, plotted on a log-scale. As expected, synonymous variants at low VAF follow the distribution predicted by genetic drift alone. Their density begins to fall-off exponentially at VAF*>* 0.03% which, combined with the age of simulated individuals of 70 years, correctly recovers the true *Nτ* = 10^5^ years used in the simulations (Figure 1D, grey curve).

#### Synonymous variants at high VAF are mostly passengers

Synonymous variants detected at VAF *> ϕ* are unlikely to be caused by drift alone. Synonymous variants reaching these higher VAFs are either passenger mutations driven to high frequency by genetic hitchhiking (Figure 1C cases II and III), or, they are synonymous variants that occurred during the earliest stages of the development of the tissue (Figure 1C case IV). Synonymous passenger mutations can occur in two distinct ways: either a clone with a ‘driver’ mutation subsequently acquires a synonymous mutation (Figure 1C case II), or, a clone with a synonymous mutation subsequently acquires a ‘driver’ mutation (Figure 1C case III). For the data and VAFs considered in this study, case II accounts for the vast majority of passengers that occur, while case III makes only a small contribution (Supplementary note 1C and Figure S2). Mutations that occur during early development (case IV) may be observed at high VAFs but for the two tissues examined in this study, estimates of developmental mutation rates suggest that they account for *<* 5% of the high VAF synonymous variants observed (Supplementary notes 1B, 3G, 5D). The density of synonymous VAFs that results from hitchhiking can be understood by first considering the case where all ‘driver’ mutations confer the same fitness advantage, *s*, and occur at a total rate, *µ*_*b*_. In this case ‘driver’ mutations that are exponentially expanding are unlikely to reach VAFs *> ϕ*_*s*_ = exp(*st*)*/*2*N τs*. The distribution of synonymous variants below *ϕ*_*s*_ has the following form

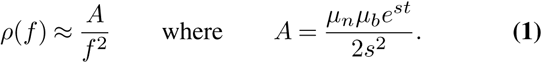

This scaling relation has two important features. First, the density of synonymous passenger mutations decays with the inverse square of the VAF (Figure 1D, orange line). Because this decay is much slower than the exponential decay of neutral clones, high VAF synonymous variants are much more likely to be passengers. Second, the amplitude, *A*, depends on the total rate of positive selection *µ*_*b*_. Therefore, provided an estimate of *s*, the amplitude of the inverse-square relationship provides an estimate for the total rate of positive selection, *µ*_*b*_. We validated this with stochastic simulations over a wide range of fitness effects and mutation rates (Supplementary note 1 and 2). The density of high VAF synonymous variants does indeed decay with the inverse square of the VAF (Figure 1D) and the amplitude of this decay accurately recovers the total rate of positive selection (Figure 1D inset).

#### A range of fitness effects and ages

When ‘driver’ mutations do not always confer the same fitness effect due to different mutation types, the expression for the density of synonymous variants from Equation (1) becomes more cumbersome and there is no longer a universal scaling with VAF due to the range of different *s*’s (Supplementary note 3F). The range of *s* that contribute most to the passenger spectrum is determined by the age of the individual as well as the shape of the DFE. At early ages, ‘drivers’ with small fitness effects contribute substantially due to their higher rate of occurrence whereas at late ages, ‘drivers’ with large fitness effects, even though they are rarer, contribute most because of their exponentially larger clone size (Supplementary Figure S9). For the typical range of ages considered in the data that follows (50-80 years) fitness effects between 9 − 15% contribute most to the passenger spectrum which falls off slightly faster than 1*/f* ^2^ (Supplementary Figure S9). Thus, even with a range of different fitness effects and a range of ages, the amplitude of high VAF synonymous variants enables estimation of the total rate of positive selection (Supplementary note 1, Equation 12).

### Many unobserved driver mutations in healthy blood

Applying our method for estimating the total rate of positive selection to ∼ 1500 synonymous variants detected in peripheral blood from Bolton et al. ^14^ and Razavi et al. ^15^ with combined total of 637 people we see striking agreement between the data and our predictions based on a model of stem cell dynamics with genetic hitchhiking (Figure 2). Most synonymous variants from ultra-deep sequencing in ^15^ are concentrated at low VAFs (*<* 0.2%) and the density of these low VAF variants decays exponentially with VAF. At higher VAFs (*>* 0.2%) we observe a clear transition whereby the decay in the density of synonymous variants begins to fall off more slowly, declining approximately with the inverse square of VAF (Figure 2A). The exponential decline in density of the low VAF synonymous variants is consistent with genetic drift in a population of haematopoietic stem cells (HSCs) where *Nτ* ≈ 50, 000 years (Figure 2A, dark grey line and Supplementary note 3C). This is in close agreement with recent estimates for *Nτ* derived from the VAF distributions of ‘driver’ mutations in clonal haematopoiesis data ^17^ and from the phylogenetic tree formed by HSCs from a middle-aged male ^3^. The slower decline in density of synonymous variants at higher VAFs (*>* 0.2%) is consistent with the roughly 1*/f* ^2^ scaling predicted for synonymous passengers (Figure 2A orange lines and Supplementary note 3C). While developmental mutations are expected to have a 1*/f* ^2^ scaling with VAF, estimates of developmental mutation rates from ^3^ suggests that only 5% of high VAF synonymous variants are developmental in origin.

**Fig. 2.**
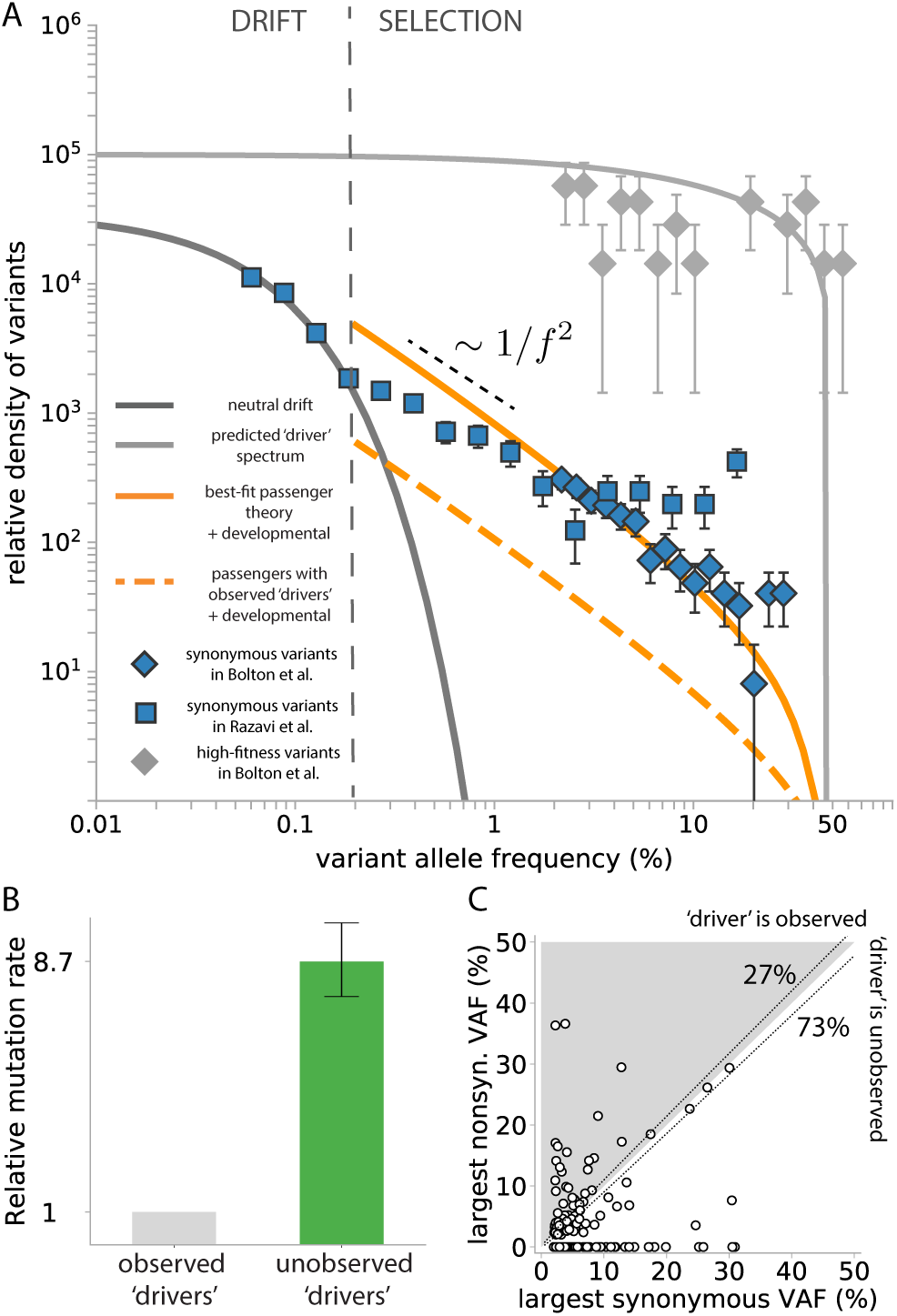
Synonymous variants in healthy blood. (**A**) The density of synonymous variants (data points) from ^14,15^ are compared to the hitchhiker prediction based on 468 cancer-associated genes (orange dashed line) and best fit (solid line). The neutral drift prediction is based on *Nτ* = 50, 000 and *µ*_*n*_ = 8.1 × 10^−4^ both inferred from Razavi et al. ^15^ (Supplementary note 3C). The density of 20 high-fitness variants (grey data points) from Bolton et al. ^14^ decline with a scaling closer to ∼ 1*/f* (flat in this log-log plot) in agreement with strong positive selection ^17^. (**B**) The rate of unobserved ‘driver’ mutation is ∼ 9-fold higher than the rate of observed ‘driver’ mutation (those occurring in the 468 cancer associated genes from ^14^). (**C**) Correlation of the highest nonsynonymous VAF and the highest synonymous VAF within the same individual in untreated patients (*n* = 200) harbouring at least one synonymous variant in Bolton et al. ^14^. Approximately 55 ± 10 individuals out of 200 (∼ 20 − 30%) harbour a putative ‘driver’ mutation detected at a higher VAF than the highest VAF synonymous variant. The diagonal dashed lines indicate the upper and lower error due to sampling noise for exactly same VAFs.

#### Synonymous VAF spectrum

To determine whether the number of synonymous passenger mutations agrees with what would be expected from the rates of observed ‘driver’ mutation, we explored two approaches. First we used estimates of mutation rates and fitness effects of 20 common high-fitness variants in clonal haematopoiesis (CH) from ^17^ to assess what fraction of high VAF synonymous variants they could explain (Supplementary note 3D). The observed density of high VAF synonymous variants is ∼ 20-fold higher than can be accounted for by hitchhiking alongside these 20 ‘driver’ mutations and suggests that ∼ 95% of positive selection is unexplained by these variants. Next, in order to assess whether this missing positive selection could be explained by mutations in other known cancer-associated genes, we inferred the DFE for putative ‘driver’ mutations using the nonsynonymous variants detected in 468 cancer-associated genes from Bolton et al. ^14^ (Supplementary note 3E). Combining this with Equation 12 we could then predict the density of synonymous passenger mutations expected based on these observed putative ‘drivers’ (Figure 2A, orange dashed line). While the scaling of this prediction with VAF is in close agreement with the data, we observe that the density of high VAF synonymous variants is ∼ 10-fold higher than can be accounted for by the DFE of these observed putative ‘driver’ mutations (solid versus dashed orange line, Figure 2A, Figure 32B). This discrepancy in amplitude, always close to 10-fold, is observed across different parameterised forms of the DFE (Supplementary note 3H). While ‘drivers’ that occur in known cancer-associated genes other than the 20 common high-fitness CH variants do indeed account for a further 5% of the missing selection, the data implies that ∼ 90% of clonal expansions in healthy blood remain unexplained and appear to be driven mutations that lie outside both the common high fitness CH variants and the 468 cancer-associated genes in the MSK-IMPACT panel ^14^.

#### Mutation co-occurrence within individuals

To test this conclusion, we considered co-occurrence of putative ‘driver’ mutations (nonsynonymous variants in the panel) with synonymous variants within the same individual (Figure 2C, Supplementary note 3I). Because most hitchhiking occurs after the expansion of ‘driver’ mutations (case II, Figure 1C, Supplementary note 1C), the VAF of synonymous passengers cannot be much larger than the VAF of the ‘driver’ mutation causing the clonal expansion (Figure 2C, shaded region). However, among individuals harbouring at least one synonymous variant from ^14^, only 20 − 30% harbour nonsynonymous variants whose VAF is higher than that of the largest synonymous variant (Figure 2C, shaded region). This therefore suggests that ∼70 − 80% of clonal expansions which drove a synonymous variant to high frequency did so without the ‘driver’ mutation being detected, implying it occurred outside the 468 cancer-associated ‘driver’ genes. This finding is in good quantitative agreement with the estimate ∼ 90% of unobserved ‘driver’ mutations inferred from the fit to the VAF distribution (Figure 2A and B).

#### Age dependence of the synonymous VAF spectrum

To further check the predictions of our model we considered the age dependence of the synonymous VAF spectrum. Our model predicts that the amplitude and shape of the synonymous VAF spectrum should exhibit strong age dependence (Supplementary Figure S9). To check this prediction, we divided 395 individuals from ^14^ into a younger age group (64 − 72 years, *n* = 194) and an older age group (*>* 72 years, *n* = 201) and plotted the synonymous VAF spectrum for each group against their respective predictions. Because mutation burden in young individuals diagnosed with cancer is possibly elevated, we excluded the youngest one-third of individuals (younger than the lower tercile at age 64). The data shows age dependence in qualitative agreement with predictions, although the dependence is not as strong as predicted by the model (Figure S15A). One possible explanation for this could be that a larger proportion of high VAF synonymous variants are developmental in origin. However because developmental mutation rate estimates (Supplementary note 3B) suggest that only 5% of high VAF synonymous variants are developmental, this weaker age dependence of the observed spectrum remains a puzzle.

### Few unobserved drivers in healthy oesophagus

The VAF distribution of ∼ 600 synonymous variants observed in healthy oesophagus from 9 individuals ^5^ can be similarly used to estimate the total rate of positive selection in healthy oesophagus. The VAF distribution of synonymous variants is again in close agreement with the near 1*/f* ^2^ scaling predicted for synonymous passengers (Figure 3A, orange line). Due to limited sensitivity at VAFs*<* 1%, no feature consistent with drift alone is apparent (Figure 3A, grey line).

**Fig. 3.**
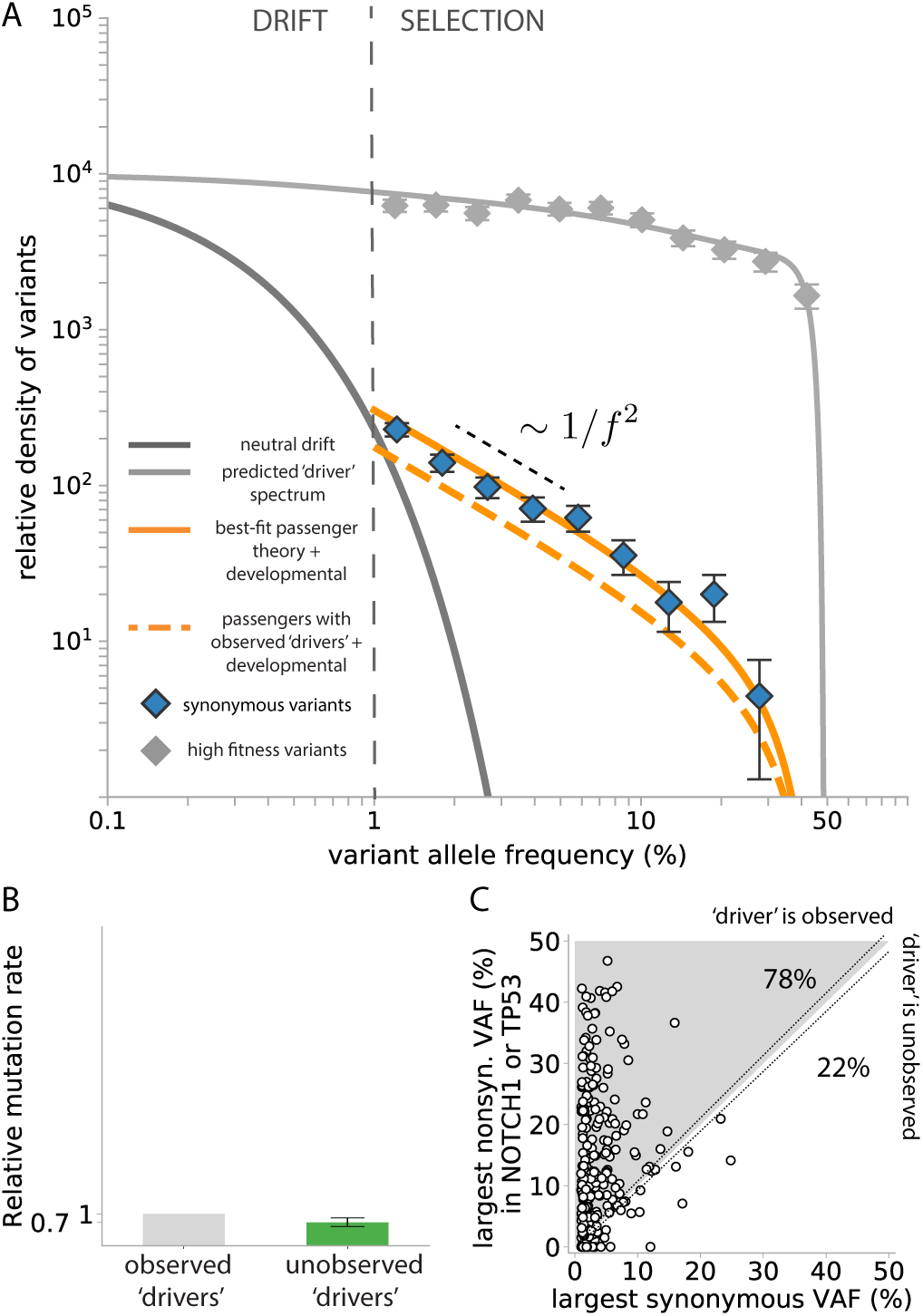
Synonymous variants in healthy oesophagus. (**A**) The density of synonymous variants (blue data points) from ^5^ are compared to the hitchhiker prediction accounting for selection from *NOTCH1* and *TP53* only (orange dashed line) and a best fit (solid line). The neutral drift prediction (dark grey line) is based on *Nτ* inferred by considering the TP53 nonsynonymous VAF spectrum (Supplementary note 5B).The density of NOTCH1 nonsynonymous variants (grey data points) decline much less rapidly with VAF, in agreement with strong selection ^17^. (**B**) The number of observed synonymous passengers is only ∼ 1.7-fold higher than can be accounted for by considering only ‘driver’ mutation in *NOTCH1* and *TP53*. (**C**) Correlation of the highest *NOTCH1* or *TP53* nonsynonymous VAF and the highest synonymous VAF within samples (*n* = 277) harbouring any detected synonymous variant. Approximately 209 − 222 individuals out of 277 (∼ 75 − 80%) harbour a putative ‘driver’ mutation detected at a higher VAF than the highest VAF synonymous variant. The diagonal dashed lines indicate the upper and lower error due to sampling noise for exactly same VAFs.

#### Synonymous VAF spectrum

To estimate the number of synonymous variants in healthy oesophagus expected due to hitchhiking with putative ‘driver’ mutations, we first estimated the fitness effects of ‘driver’ mutations (Supplementary note 5B). We estimate that nonsynonymous mutations in *NOTCH1* have a fitness effect of *s* = 11% per year and occur at a haploid rate of 3.3 × 10^−5^ per year, while those in *TP53* confer a fitness effect of *s* = 10% per year and occur at a haploid rate of 1.4 × 10^−5^ per year. There is little evidence of strong selection outside of these genes (Supplementary note 5D, ^5,18^). Using these estimates, we find that the observed density of high VAF synonymous variants is ∼ 1.7-fold higher than can be accounted for by ‘driver’ mutations in these two genes alone (Figure 3B). This suggests that *TP53* and *NOTCH1* account for more than half of the genome-wide rate of positive selection.

#### Mutation co-occurrence within individuals

To test this conclusion, we considered mutation co-occurrence of non-synonymous driver mutations in *NOTCH1* and *TP53* with synonymous variants within the same biopsy sample (Figure 3C). The synonymous VAF distribution suggests that ‘driver’ mutations in *NOTCH1* and *TP53* account for the majority of clonal expansions, therefore we predict that in a large fraction of biopsy samples the synonymous variants will be found at a lower VAF than the largest nonsynonymous variant within these two genes (Figure 3C, shaded region). Consistent with this prediction, we observe that in 75 − 80% of samples with detected synonymous variants, the largest nonsynonymous variant in *NOTCH1* or *TP53* is found at a higher VAF than the largest synonymous variant (Figure 3C, Supplementary note 5E). This analysis implies that genetic hitchhiking in healthy oesophagus is dominated by mutations in these two genes alone and that few other strong ‘driver’ mutations exist elsewhere in the genome.

#### Age dependence of the synonymous VAF spectrum

To further check the predictions of our model we again considered the age dependence of the synonymous VAF spectrum. We divided the samples into a younger age group (ages = 21, 25, 37, *n* = 273) and an older age group (ages = 57, 69, 73, *n* = 284) and plotted the synonymous VAF spectrum for each group against their respective predictions. The data shows clear age dependence in qualitative agreement with model predictions (Figure S15B). However, the observed age dependence is again slightly weaker than predicted by the model.

## Discussion

We have developed a population genetic framework to quantify the total rate of positive selection in healthy blood and oesophagus by considering the density of high VAF synonymous variants. The key intuition behind our analysis is that because synonymous mutations are likely to have been carried to high VAF by hitchhiking with ‘driver’ mutations, the number of high VAF synonymous mutations observed across multiple samples can provide an estimate for the *genomewide* rate at which ‘driver’ mutations are acquired and therefore the fraction of driver mutations missed by narrow cancer-associated gene panels.

### Blood

In blood, the majority of synonymous variants reaching high VAF do so by virtue of hitchhiking with a ‘driver’. The large number of high VAF synonymous variants, however, suggests that most clonal expansions are caused by ‘drivers’ that fall outside of cancer-associated gene panels. Consistent with this finding, most high VAF synonymous variants are not observed to co-occur with any candidate ‘driver’ mutation. The large number of unobserved ‘drivers’ likely include mCAs ^10,11,29,30^, large structural variants ^31,32^, coding mutations in non-cancer-associated genes, mutations in regulatory regions ^33–36^ and, possibly, epigenetic alterations ^37,38^. We hypothesize that a more comprehensive survey of these alterations will yield many further mutations that will be shown to be capable of driving clonal expansions in blood ^16^. To demonstrate this, we analysed the VAF spectrum of the copy number change resulting in the loss of the Y chromosome studied in ^29^. These data suggest that loss of Y occurs at a rate of ∼ 10^−6^ per cell per year and confers a strong fitness advantage of *s* ∼ 10% per year (Supplementary note 6). Using these estimates, Y loss can only account for *<* 10% of the missing positive selection in blood. While this only affects males, recent evidence that mCAs are relatively common drivers of clonal haematopoiesis ^10,11^, suggests that the collective effect of all mCAs might account for a substantial proportion of the missing selection and this is an important area for future research.

One remaining puzzle is that the data from blood shows a weaker age dependence than is predicted by the model. One possible factor contributing to this lack of quantitative agreement is that the cohort in ^14^, on which our age analysis is based, are treatment-naive patients with at least one solid cancer. This cohort, especially those diagnosed young, therefore may not reflect completely healthy haematopoiesis.

### Oesophagus

In oesophagus, the picture is quite different. While high VAF synonymous variants are also consistent with being caused by hitchhiking, our analysis shows that the number of high VAF synonymous variants is broadly what would be expected if the only mutations capable of driving clonal expansions were those in *NOTCH1* and *TP53*. In support of this, most high VAF synonymous variants in oesophagus co-occur with a larger non-synonymous variant in either *NOTCH1* or *TP53*. We hypothesize that strong ‘drivers’ of clonal expansions in normal oesophagus residing elsewhere in the genome are therefore rare. One prediction made by this hypothesis is that, if esophageal squamous-cell carcinomas (ESSC) begins with a clonal expansion in normal tissue, mutations in one of these two genes should be an early event in almost all ESSCs. Mutations in these two genes are indeed observed in ∼ 90% of ESSCs and, at least in the case of *TP53*, are estimated to be early events ^39,40^.

### Generalisation to other tissues

Our framework for estimating the total rate at which clonal expansions occur in healthy tissues is general and may be applicable to other tissues where the growth of ‘driver’ mutations is unhindered by tissue structure (e.g. skin). However our model will require further development to be applied in settings where there is strong tissue structure (e.g. colonic crypts or lung). Furthermore, our model relies on a number of assumptions, outlined below, that merit careful consideration.

### Fitness effects of ‘driver’ mutations

The amplitude of high VAF synonymous variants depends exponentially on the fitness effects of linked ‘driver’ mutations. Therefore, uncertainties in the distribution of fitness effects (DFE) of driver mutations can lead to large uncertainties in the estimates for total ‘driver’ mutation burden. To investigate the robustness of our inferences to these effects we explored a number of different possible forms for the DFE (Supplementary note 3H). While the details of the functional forms for the DFE naturally make small quantitative differences to our results they do not alter the broad qualitative conclusions.

### Developmental mutations

The contribution of mutations that arise during early development (Figure 1) to the synonymous VAF spectra relies on estimates of mutation rates per cell division during the development of the tissue. In blood, where robust estimates of the developmental mutation rates can be obtained from single-cell-derived whole genome sequencing of HSCs (^3^ and Supplementary note 3B), the contribution of developmental mutations is estimated to be ∼ 5% (Figure S10). In oesophagus, we are not aware of any robust estimates of developmental mutation rates and therefore we used the estimated developmental mutation rates from blood as a starting point. The consistency between the observed amplitude of high VAF synonymous variants and predicted amplitude based on mutations in *NOTCH1* and *TP53* suggests that the contribution from development is also small.

### Synonymous mutations subject to selection

An important assumption in our analysis is that synonymous variants are selectively neutral during somatic evolution. Evidence from cancer genomes suggests that some synonymous variants have functional consequences ^41,42^, possibly due to codon usage bias ^43,44^. However, it is likely these variants constitute a small minority of all synonymous variants and confer weak fitness effects ^36^.

### A simplified branching model

We have used an intentionally simplified model of stem-cell dynamics in which many complicating factors such as ageing, the micro-environment and spatial effects are not explicitly accounted for. While these effects likely do influence stem-cell dynamics, in previous work we have shown that this simple stochastic branching model can explain many of the quantitative aspects of the VAF data ^17^. Moreover two features in the data here suggest this simple model captures much of the relevant behaviour. First, the data clearly shows that variants under strong positive selection have a different VAF dependence from passenger mutations, scaling as 1*/f* versus ∼ 1*/f* ^2^ as the model predicts. Second, the blood sequencing data clearly shows a transition at ∼0.2% VAF from drift-dominated to selection-dominated behavior that quantitatively agrees with the predictions of this model and with previous estimates of *Nτ* (Figure S6, ^3,17^). This latter point demonstrates the value of studies with sensitive sequencing that are capable of detecting neutral clones ^7,15,45^ as they provide important internal consistency checks on the model. The limited quantitative agreement for age-dependence of the synonymous VAF spectra however points to limitations of this simplified theory (Figure S15).

### Conclusions

We have shown that synonymous variants that reach high VAF via genetic hitchhiking provide an important self-consistency check on the number of mutations driving clonal expansions. This self-consistency, or lack thereof, provides information on the genome-wide rate of positive selection including contributions from alterations hard to detect or not often looked for, such as mCAs, SVs, non-coding, and epigenetic changes. In blood (and likely in other tissues too) it appears that SNVs are only the tip of the ‘driver’ mutation iceberg and this is evidenced by recent studies showing the prevalence of mCAs that cause large clonal expansions in healthy blood ^10,11^. As somatic evolution in various tissues becomes more comprehensively mapped, many more drivers of clonal expansions will emerge, leading to better understanding of cancer evolution that informs treatment decisions and points towards potential therapeutic targets.

## Acknowledgements

We thank Kelly Bolton, Ahmet Zehir, Elli Papaemmanuil for sharing unpublished data. We also thank David Solit, Pedram Razavi and Jorge Reis-Filho for sharing data and Inigo Martincorena for sharing data and for helpful discussions. Y.P.G. P, C.J.W and J.R.B are funded by the CRUK Cambridge Centre and CRUK Early Detection Programme. J.R.B is supported by a UKRI Future Leaders Fellowship. D.S.F and J.R.B. are supported by the Stand Up to Cancer Foundation and the National Science Foundation via PHY-1545840.

## Data and code availability

All code used in this study will be available on the Blundell lab Github page.

## Supplementary Note 1: Derivation of synonymous variant allele frequency spectrum

### A. Neutral drift

During homeostasis, mutations can occur in a single cell, creating a single mutant lineage that can drift, expand or die out. If the population size of stem cells is constant, the feeding rate *Nτµ*_*n*_ (*τ* is the time in years between symmetric stem cell divisions) to neutral mutant lineages is also constant. The resulting variant allele frequency spectrum for non-extinct lineages is

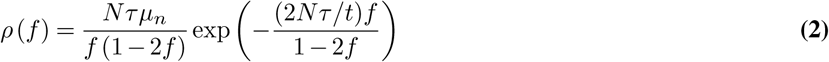

where *µ*_*n*_, *t* are measured in years and *τ* is the symmetric stem cell division time in units of years. After performing logarithmic transform *l* = log(*f*), the neutral drift frequency spectrum is essentially flat for small *f* because of the 1*/f* scaling. The spectrum exponentially decays at *f > ϕ* = *t/*2*Nτ*. The above derivation assumes no clonal interference among lineages, which is true for neutral mutant lineages remaining small under drift.

This predicts that there will be a high density of synonymous mutations confined to low VAF because they are under the influence of drift and experience no selection. This feature is clearly observed in simulated data (Figure S3) as well as in the low VAF part of the data from healthy blood with an amplitude that agrees with estimates of *Nτ* from literature ^3,17^ (Figure S6).

#### Stochastic simulations

We used stochastic simulations to validate our predictions and test the robustness of our model. In our simulations, a cell gives birth to a number of offspring in an interval of time *δt*, which is randomly distributed according to the Poisson distribution with expected value *Bδt* but also produces a number of cell deaths following the Poisson distribution with expected value *Dδt. B* and *D* represent birth rate and death rates. Using units of time which are generations (i.e. *D* = 1), the simulation is run at intervals of *δt* = 0.1, i.e. one-tenth of a generation as an approximation to a continuous time process. Mutation fitness effects are incorporated in birth rates *B* = 1 + *s*_rel_*τ*, where *s*_rel_ is the relative fitness to the mean fitness of the population per year, which typically grows in time as more and more ‘driver’ mutations are introduced into the population, and *τ* is the generation time measured in years. The probability of producing a new mutant cell in the next generation from an existing clone of size *n* is always *u n*, where *u* represents the mutation rate per generation. We assume fitness effects of new mutations are additive and always non-negative in their absolute values such that a clone with two mutation with fitness effects *s*_1_ and *s*_2_ in a wild-type population would have a fitness advantage of *s*_1_ + *s*_2_.

The simulated neutral VAF spectra matches our predictions for neutral drift and for passengers (Figure S1). A passenger refers to any neutral mutation that coexists with at least one ‘driver’ mutation. Simulated VAF reflects diploid cells, i.e. VAF = 0.5 is the maximum corresponding to a 100% cell fraction. Our theory also correctly accounts for passengers arising via two different routes of genetic hitchhiking (Figure S2).

### B. Developmental mutations

Another possible way that neutral synonymous mutations can reach high frequencies is if they occur during the early stages of development. Humans develop from a single cell into a full organism via many cycles of cell divisions. It stands to reason that mutations occurring during the early stages of the development of the blood system, will be present at high frequencies in the adult HSC population. We assume early embryonic growth is deterministic with high division rates, thus can be accurately modelled by exponential growth. Assuming the entire HSC population in a person grows as 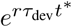 with time *t*^*^ measured in years from a single ancestor, where *r* is the growth rate per time and *τ*_dev_ is the division time during development measured in years. Given the developmental mutation rate per year 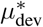, the probability a synonymous mutation enters at time *t*^*^ is 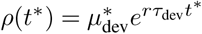 per year. Under the deterministic assumption and assuming only birth, the neutral mutant lineage that entered grows as 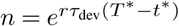 for 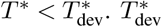 marks the end of the developmental stage. By applying this transformation between *n* and *t*^*^ to *ρ*(*t*^*^), the clone size distribution of developmental mutations is

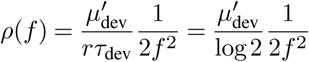

where the parameter combination *rτ*_dev_ has been replaced by log 2 in a birth-only scenario. The VAF *f* in a diploid cell population is related to *n* via 2*f* = *n/N*. The distribution does not depend on the growth rate *r* or the age of the person and only depends on 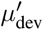 which is the mutation rate per cell per cell doubling. To validate that our theory correctly captures the density of developmental synonymous mutations, we ran stochastic simulations and confirmed that the density of synonymous mutations that are developmental in origin matches with our theoretical predictions (Figure S3).

#### Simulating developmental stage

We also set up simulations to reflect the more realistic scenario where the stem cell population grows from a single cell during development before reaching homeostasis. During development, the stem cell population grows deterministically at an exponential rate 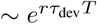, where *T* is time measured in years. Mutations enter according to a distribution of fitness effects. All mutations are treated as neutral and expand deterministically during development until the entire stem cell population reaches the adult population size, at which point stem cells’ fates become stochastic and cells harbouring beneficial ‘drivers’ are likely to produce more offspring.

Our theoretical prediction (Supplementary note 1B) for high VAF developmental mutations matches with the simulated VAF spectra of neutral mutations that arise during development (Figure S3, green diamonds). The simulated VAF spectrum has a characteristic 1*/f* ^2^ scaling and its amplitude at high-VAF is consistent with model predictions (Figure S3, green line). The simulated neutral VAF spectrum (Figure S3, blue diamonds) can be accounted for by effects of neutral drift at low frequencies (Figure S3, grey line, Supplementary note 1A), and contribution from genetic hitchhiking and development (Figure S3, blue line, Supplementary note 1C) at intermediate and high VAFs.

Developmental mutations may occur that can later act as ‘drivers’ to cause expansions during homeostasis after the clone acquires further mutations. The contribution of these ‘driver’ mutations to those arising during homeostasis depends on the developmental mutation rates and their mutation fitness. The additional fold to the passenger spectrum contributed by these developmental ‘driver’ mutations is approximately equal to the probability of a ‘driver’ mutation arising in a population, i.e. 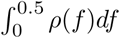, which is negligible.

### C. Passengers

Genetic hitchhiking occurs when a neutral passenger mutation co-occurs in the same cell with a ‘driver’ mutation under positive selection and is brought to high frequencies in the cell population as a result. This can occur via two distinct routes - either the neutral mutation hitchhiked on a ‘driver’ clone already in existence (termed *beneficial-first route*) or a neutral clone survives drift for long enough until it gains a ‘driver’ mutation (termed *neutral-first route*). As derived more formally below, neutral lineages grow slowly and are unlikely to survive for long periods of time until a subsequent ‘driver’ mutation occurs, thus most passengers arise by hitchhiking with a ‘driver’ clone that already existed.

#### Passengers via beneficial-first route

The wild-type stem cell population feeds ‘driver’ mutations at rate *τµ*_*b*_ per generation, which results in the following clone size distribution for beneficial ‘driver’ mutants at time *t*′ years:

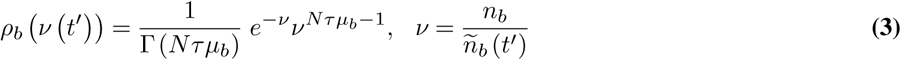

For ‘driver’ mutants with fitness *s* per year, *ñ*_*b*_ (*t*′) = (*e*^*st*′^− 1)*/sτ*. Passengers can enter the beneficial clone at any time *t*′ and establish, after which the passenger lineage grows exponentially like any other beneficial ‘driver’ lineage. Because neutral mutation rate is low, a passenger enters with a probability of approximately *τµ*_*n*_*n*_*b*_(*t*′) per generation at all times, only determined by the expected size *ñ*_*b*_(*t*′) of the beneficial clone at time *t*′, which is under exponential growth. The probability of a passenger entering at *t*′ is

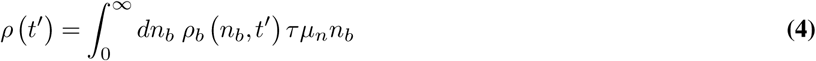

which integrates to

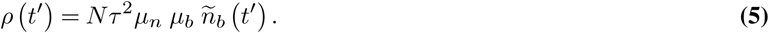

where *ñ*_*b*_ (*t*′) = (*e*^*st′*^− 1)*/sτ*. To account for the effect of ‘driver’ mutants sweeping in the stem cell population, the feeding rate and growth rate of passengers hitchhiking with a particular ‘driver’ are stopped once the expected clone size of this particular ‘driver’ is large enough that the relative fitness to the stem cell population of another same ‘driver’ mutation arising becomes less than 5% per year.

Based on the exact form of the clone size distribution of a non-extinct single mutant (not conditioned on survival: note that this is also why we do not have to consider the establishment probability explicitly) ^22^,

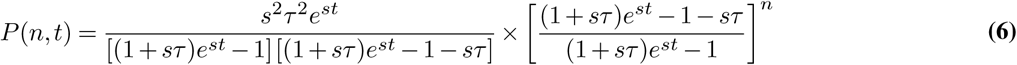

we use the approximated form

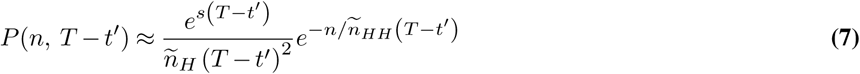

for non-extinct lineages and integrate with Equation 5 for mutants entering at time 0 *< t*′ *< T* where the expected driver clone size *ñ*_*H*_ (*T* − *t*′) is approximated to be *e*^*s*(*T*−*t*′)^*/sτ*. The contribution to the total passenger population by neutral mutations that hitchhiked a beneficial ‘driver’ clone (Figure 1A, case II) is then characterized by the clone size distribution

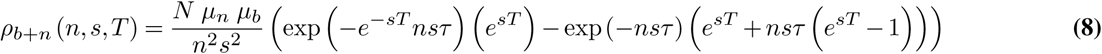

where *T, s* are measured in units of years and *τ* is the symmetric stem cell division time in units of years. We validated this expression against simulated data (middle column, Figure S2).

#### Passengers via neutral-first route

If neutral lineages survive and drift to moderately high frequencies, a small number of them will acquire a subsequent beneficial ‘driver’ mutation and become passengers (Figure 1A, case III). This occurs when a beneficial ‘driver’ mutation arises within neutral lineages *n*_*n*_ following the following statistics at time *t*′ years

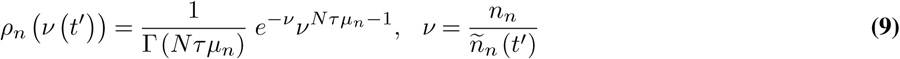

where the expected size *ñ*_*n*_(*t*′) of the neutral lineage is growing only linearly as *ñ*_*n*_(*t*′) = *t*′*/τ*.

Following the same derivation steps and approximating *ñ*_*HH*_ ≈ *e*^*s*(*T*−*t*′)^*/sτ*, the contribution to the total passenger population of neutral passengers arising via neutral mutants acquiring a further beneficial ‘driver’ (Figure 1A, case III) is then characterized by the clone size distribution

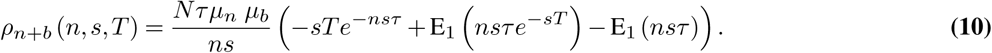

We again validated this expression against simulated data (rightmost column, Figure S2).

The full passenger spectrum *ρ*_HH_ (*f, s, T*) caused by ‘driver’ mutations with fitness *s* is the sum of formula 8 and formula 10 and transformed to variant allele frequency *f* via 2*f* = *n/* (*n* + *N*).

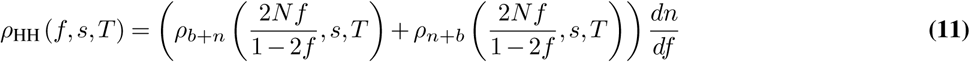

### The synonymous VAF spectrum

Therefore, the spectrum of neutral mutations consists of three components: neutral mutants under drift, developmental mutations and passengers (Figure S3).

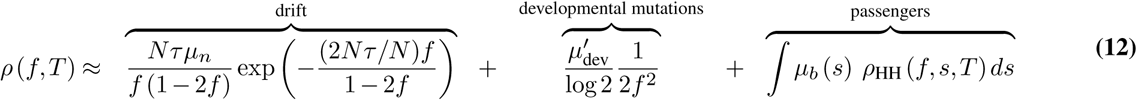

noting *T* is the age of the person, 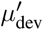 is the mutation rate per cell per cell doubling during development and *µ*_*b*_(*s*) represents the distribution of fitness effects (DFE) of ‘driver’ mutations that cause clonal expansions. Neutral and beneficial ‘driver’ mutation rates *µ*_*n*_ and *µ*_*b*_ are both measured per year. The DFE for each tissue can be inferred using the VAF spectra of ‘drivers’ and its form determines the shape of the passenger VAF spectrum.

**Fig. S1.**
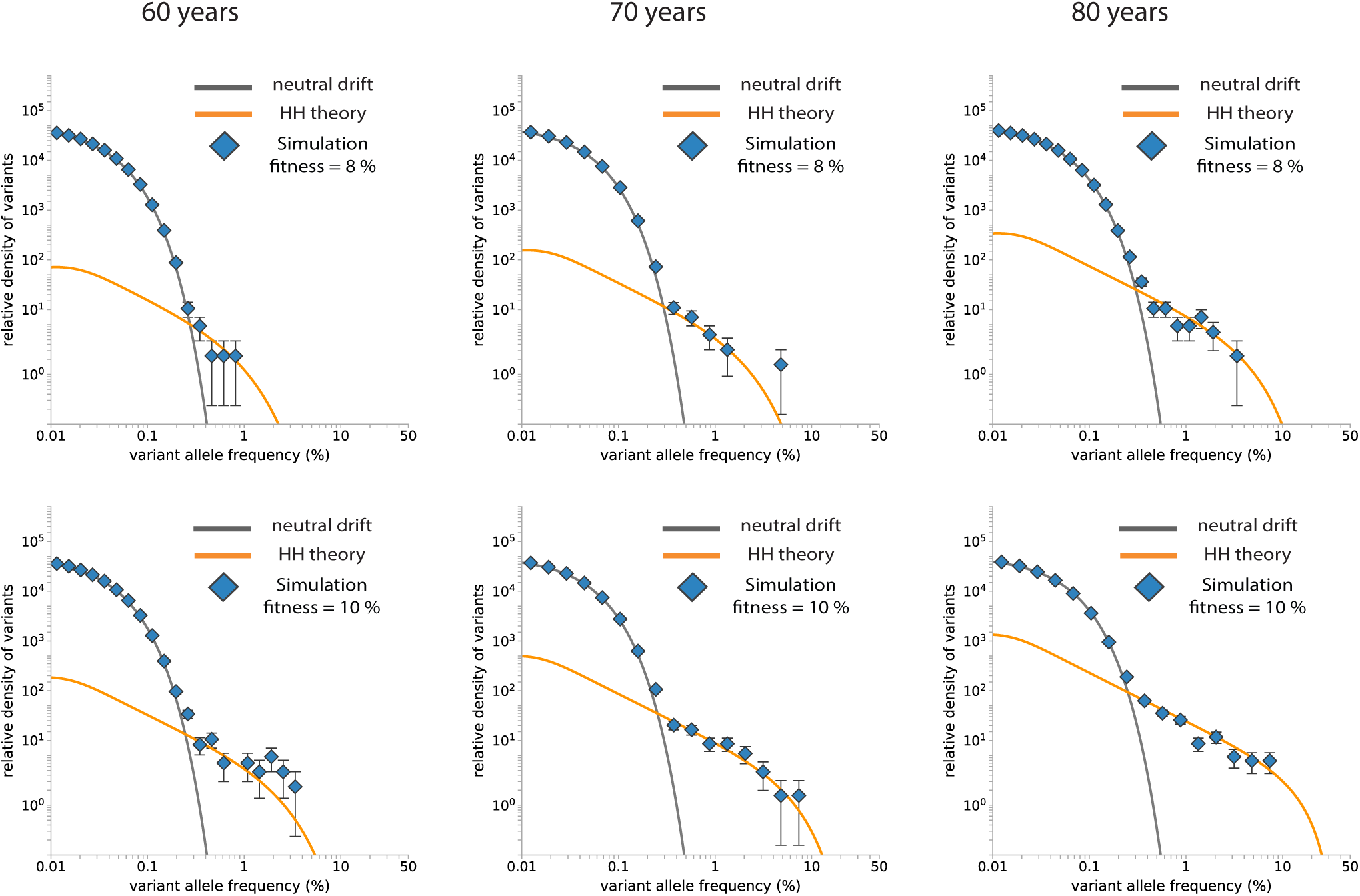
Validation of neutral drift. The neutral drift prediction (dark grey line) VAF spectra matches with the distribution of low-frequency neutral mutations in simulation (blue diamonds) at different times (*T* = 60, 70, 80 years for *τ* = 1 year, *µ*_*b*_ = 10^−5^, *µ*_*n*_ = 10^−4^, *N* = 10^5^) The passenger prediction (orange line) matches with the distribution of high-frequency neutral mutations (blue diamonds) at different selection levels (row 1 corresponds to *sτ* = 0.08 and row 2 corresponds to *sτ* = 0.1, for *τ* = 1 year). The passenger VAF spectrum is more age-dependent than the neutral drift VAF spectrum. For each set of parameters, 15000 simulations were performed for each set of parameters.

**Fig. S2.**
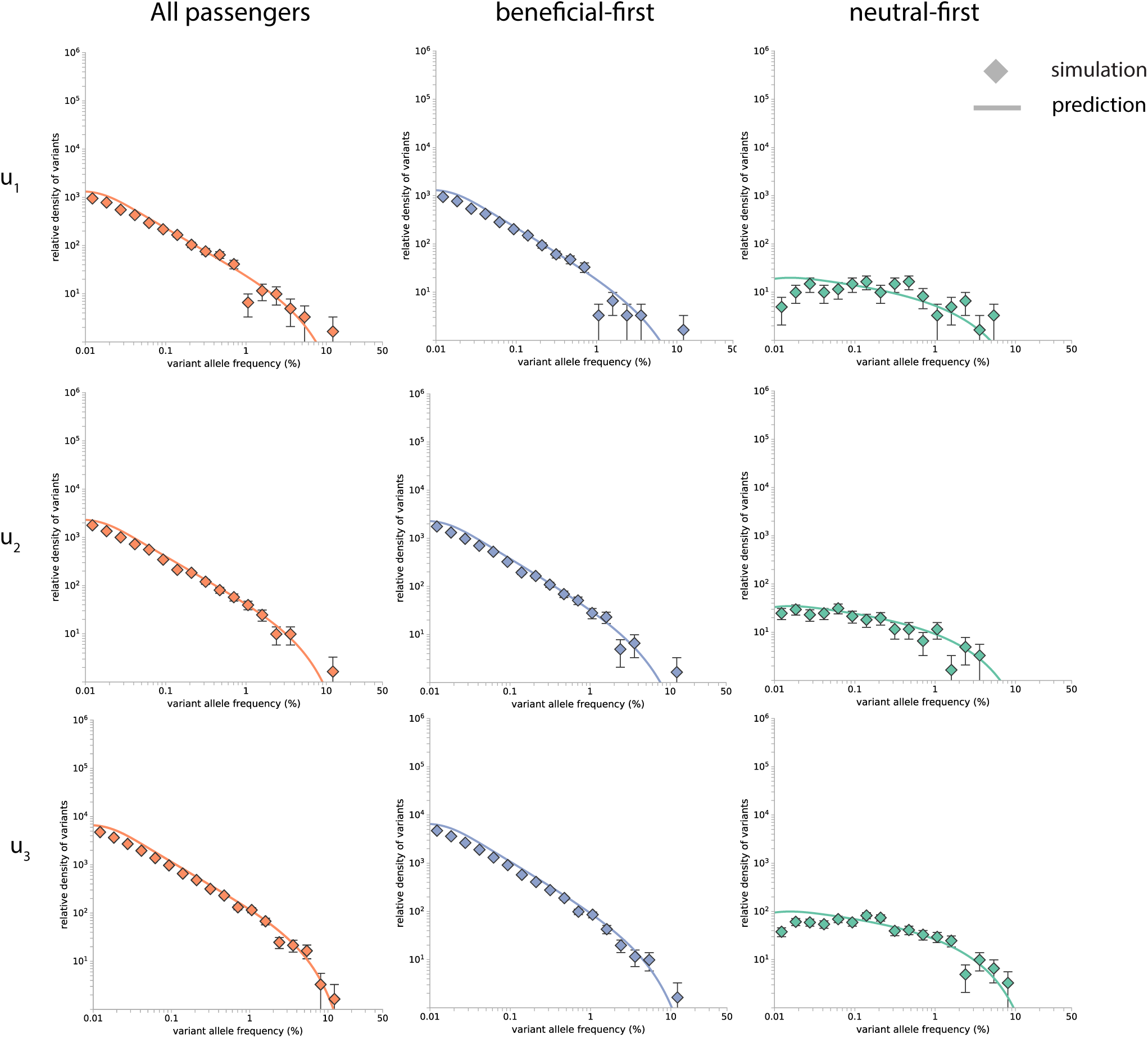
Validation of two classes of passengers. Predicted VAF spectra for passengers via ‘beneficial-first’ and ‘neutral-first’ route match with simulations. There are 15000 simulation runs corresponding to *µ*_*b*_ = *u*_1_ = 4 × 10^−6^, *µ*_*b*_ = *u*_2_ = 7 × 10^−6^, and *µ*_*b*_ = *u*_3_ = 2 × 10^−5^ respectively, all with the same neutral mutation rate *µ*_*n*_ = 10^−4^, *sτ* = 0.1, *N* = 10^5^, and examined at *T* = 70 for *τ* = 1 year. Their total in the simulations also matches with the prediction of the total passenger VAF spectra.

**Fig. S3.**
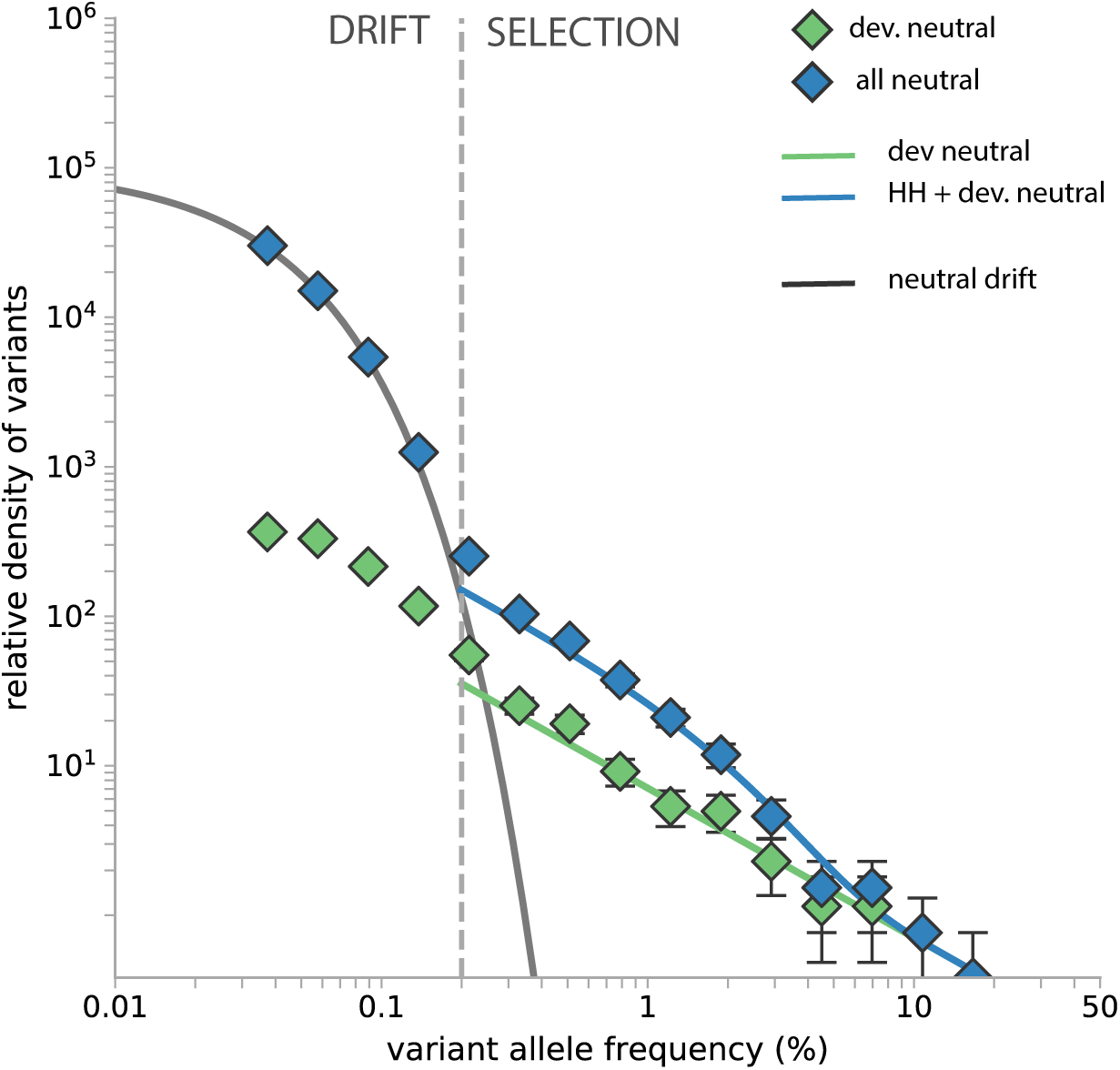
Validation of developmental theory. Our theoretical prediction (Supplementary note 1B) for high VAF developmental mutations matches with the simulated VAF spectra of neutral mutations that arise during development (green diamonds). The VAF spectrum of developmental mutations is a 1*/f* ^2^ relation. Its amplitude depends on developmental mutation rates and is not age-dependent. The simulated neutral VAF spectrum (blue diamonds) can be accounted for by effects of neutral drift at low frequencies (grey curve), and contribution from genetic hitchhiking and development (blue curve) above the observed drift limit Ψ = 2 × 10^−3^, i.e. the VAF at which the abundance of mutants driven by drift become less than that of hitchhiking mutants (Supplementary note 3C, vertical grey dashed line). (parameters: *r* = 1.2, *T* = 70 for *τ* = 1 year, *N* = 10^5^, *sτ* = 0.1; beneficial and neutral mutation rates *u*_*b*_ = 1.7 × 10^−6^ and *u*_*n*_ = 0.00010 per generation during development; *u*_*b*_ = 1 × 10^−5^ and *u*_*n*_ = 0.0006 per generation during homeostasis).

## Supplementary Note 2: Recovery of ‘driver’ mutation rate and selection level in simulations

We tested the ability of our model to accurately recover the total ‘driver’ mutation rate and the ‘driver’ mutation fitness across a range of mutation rates. First a two-parameter maximum-likelihood fit for fitness effect and mutation rate was performed on the ‘nonsynonymous’ VAF spectrum generated in simulations following the same procedure as outlined in ^17^. Second, the inferred fitness effect was then fixed at its maximum-likelihood value and a one-parameter fit was then performed on the ‘synonymous’ VAF spectrum for the total ‘driver’ mutation rate. Optimization of the likelihood was performed with the ‘Nelder-Mead’ algorithm across bins in log-scale with respect to reverse cumulative densities above the observed drift limit Ψ ≈3 × 10^−3^, i.e. the VAF at which the abundance of mutants driven by drift become less than that of hitchhiking mutants (Supplementary note 3C), and below the maximum VAF any variant under consideration attains, beyond which the detectability of variants is limited by the number of simulation runs. This approach was able to accurately recover total ‘driver’ mutation rates (Supplementary Figure S4). The modest but systematic underestimate of the total rate for high ‘driver’ mutation rates was due to clonal interference between clones within an individual. This effect is not considered here because clonal interference between two or more high VAF clones is uncommon in both the blood and the oesophagus data. However it becomes important when *Nτµ >* 1.

**Fig. S4.**
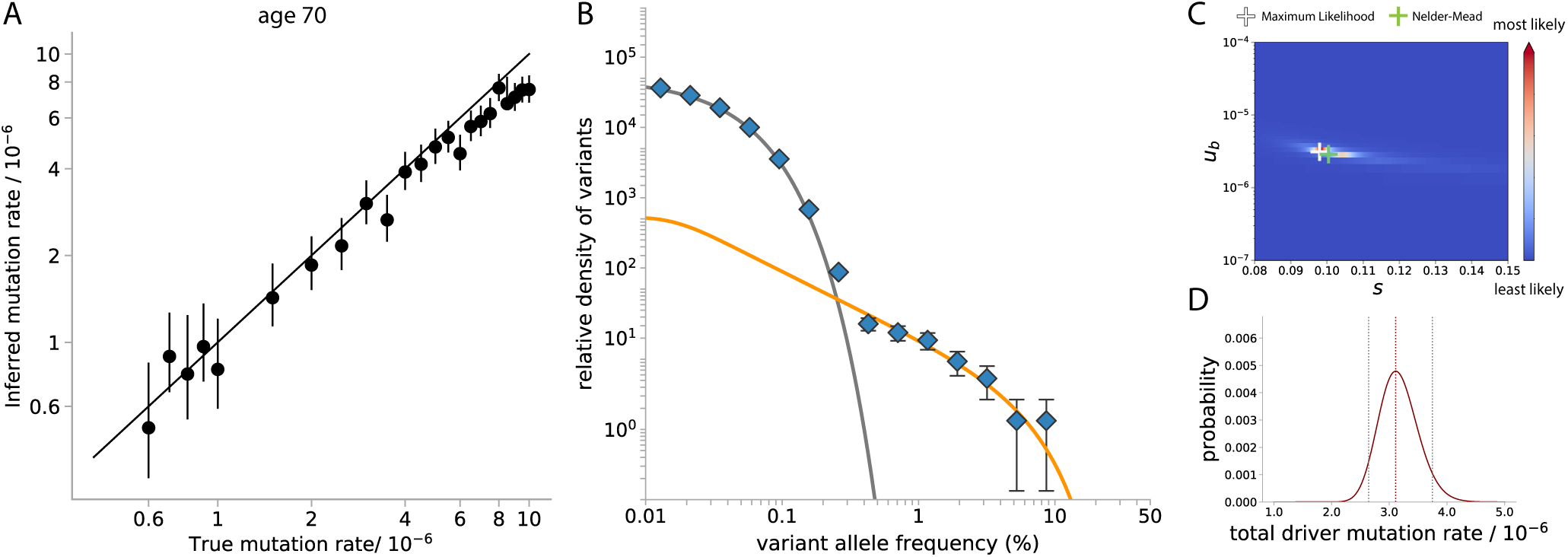
Model performance in recovering driver mutation rates in simulations. (A) Our method is able to recover ‘driver’ mutation rates accurately across a range of mutation rates. At higher ‘driver’ mutation rate, it is mainly limited by clonal interference which causes clones to reach sizes lower than that predicted by our theory. (B) An instance of the simulation run at ‘driver’ mutation rate *µ*_*b*_ = 3 × 10^−6^ (*τ* = 1 year). The neutral mutation frequency spectrum above Ψ = 3 × 10^−3^ (dotted vertical line) is fitted with our passenger prediction to infer the underlying ‘driver’ mutation rates driving the expansions. (C) The likelihood plot shows the fit for the ‘driver’ mutation rate and fitness by examining the ‘nonsynonymous’ variant allele (i.e. ‘driver’ mutation) frequency spectrum only. It is overlaid with the maximum likelihood value (white cross) and best-fit value found by the Nelder-Mead optimization algorithm (green cross). (D) The likelihood plot shows the best-fit value as well as 95% confidence intervals for the inferred total ‘driver’ mutation rate from the ‘synonymous’ variant allele (neutral mutation) frequency spectrum based on the inferred fitness.

## Supplementary Note 3: Synonymous variants in healthy blood

### A. Data processing

Two studies ^14,15^ reported synonymous variants in healthy blood using the MSK-IMPACT panel which targets a range of cancer-related driver genes. Because we are interested in the progression of clonal evolution in healthy blood, only individuals without haematologic malignancies or precursor conditions were included. Individuals that had received cancer treatment were also excluded as chemotherapy/radiotherapy has been shown to alter the selective environment within the body ^46^. Bolton et al. ^14^ involves 590 untreated patients with non-haematological cancers. They were sequenced to depths that enabled us to probe the shape of the synonymous VAF distribution at *f >* Ψ which is the VAF at which the abundance of mutants driven by drift become less than that of hitchhiking mutants and where hitchhikers are expected to dominate. On the other hand, Razavi et al. ^15^, although smaller with only 47 controls, had a greater sensitivity that enabled us to probe the shape of the synonymous VAF distribution in the *f <* Ψ regime where neutral drift is expected to dominate (i.e. *f < ϕ* for participants’ ages in the cohort) (Supplementary note 3C).

In both studies, the MSK-IMPACT panel which covers a varying number of genes (341, 410, 468 or 508) was used to achieve detection limits as low as 0.013% (Razavi et al. ^15^) and 2% (Bolton et al. ^14^). We trimmed variant data in Razavi et al. ^15^ between 0.05% and 20% to avoid false negatives at low frequencies impacting analyses and excluded germline variants and other mismapping artefacts at high frequencies (1297 variants were analysed). We did not trim data in Bolton et al. ^14^ which lie between 2% and 50% VAFs (221 variants were analysed). Because studies vary in the number of individuals, sequencing depth and sequencing panel size, to compare data across studies, we first binned the variants by VAF and divided through by the number of individuals in each study to obtain a density. We then normalized these densities by *µ*_*n*_, the study-specific synonymous mutation rate across the panel used in that given study. The mutation rates were calculated based on nucleotide context (as in our previous work ^17^) which was based on the whole-genome sequencing data presented in another study ^3^. Data in Bolton et al. ^14^ was normalized based on the FDA-approved MSK-IMPACT, a 468-gene tumor sequencing assay cancer panel which covers approximately 1.5 Mb of the genome. Because part of Bolton et al. ^14^ data was sequenced on older versions of MSK-IMPACT which included fewer genes, the normalized VAF densities are a lower bound to actual densities. Our predicted number of unseen ‘drivers’ is therefore a lower estimate. Data in Razavi et al. ^15^ was normalized based on a genomic region size of 2.13 Mb, quoted in the original paper ^15^.

### B. Developmental mutations

The VAF spectrum of synonymous mutations from healthy blood scales roughly as ∼ 1*/f* ^2^ at high VAF (Figure S5) which is consistent with our theoretical predictions for both genetic hitchhikers and possible developmental mutations (Equation 12). How many of the high VAF synonymous mutations are likely developmental in origin? The contribution of developmental mutations can be calculated using Equation 12 where 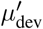 is the developmental mutation rate which we infer by applying our developmental theory (Supplementary note 1B) and fitting to whole-genome sequencing data obtained from a 59 year-old man with no known myeloid driver mutations detected ^3^ assuming the selected cell colonies are an unbiased representation of the stem cell population of the person.

Estimates obtained here for developmental mutation rates are higher than the the diploid rate of 1.2 mutations per cell per cell doubling for the first few cell doublings quoted in ^3^ because our method averages over a longer time (i.e. considers the density of mutations at lower VAFs). Because of contributions from neutral drift and hitchhiking effects during somatic evolution, the inferred developmental mutation rate is an upper bound to the actual average mutation rate before the stem cell population reaches homeostasis. The threshold *f*′ is therefore chosen to minimize these effects yet allows estimation of the developmental mutation rate averaged over the period of growth that is relevant to intermediate to high VAF mutations. Because we assume only cell birth during development, the developmental mutation rate per doubling, i.e. in units of time that the stem cell population doubles in size, can be estimated. Using this approach, the average developmental mutation rate per cell per doubling is ≈ 3.7 (95% CI interval: 3.3 − 4.2) and is applied to analyses in healthy blood and healthy oesophagus (Supplementary note 3, 5D).

The contribution of developmental synonymous mutations that results from this mutations rate has been accounted for in our analysis based on the ages of the individuals ^5,14,15^ and its contribution is small relative to the genetic hitchhikers (Figure S10A and S17A, green line). From the relative amplitudes of the developmental prediction and the data, we estimate that *<* 5% of high VAF synonymous mutations in healthy blood are developmental in origin and thus that the majority are genetic hitchhikers.

**Fig. S5.**
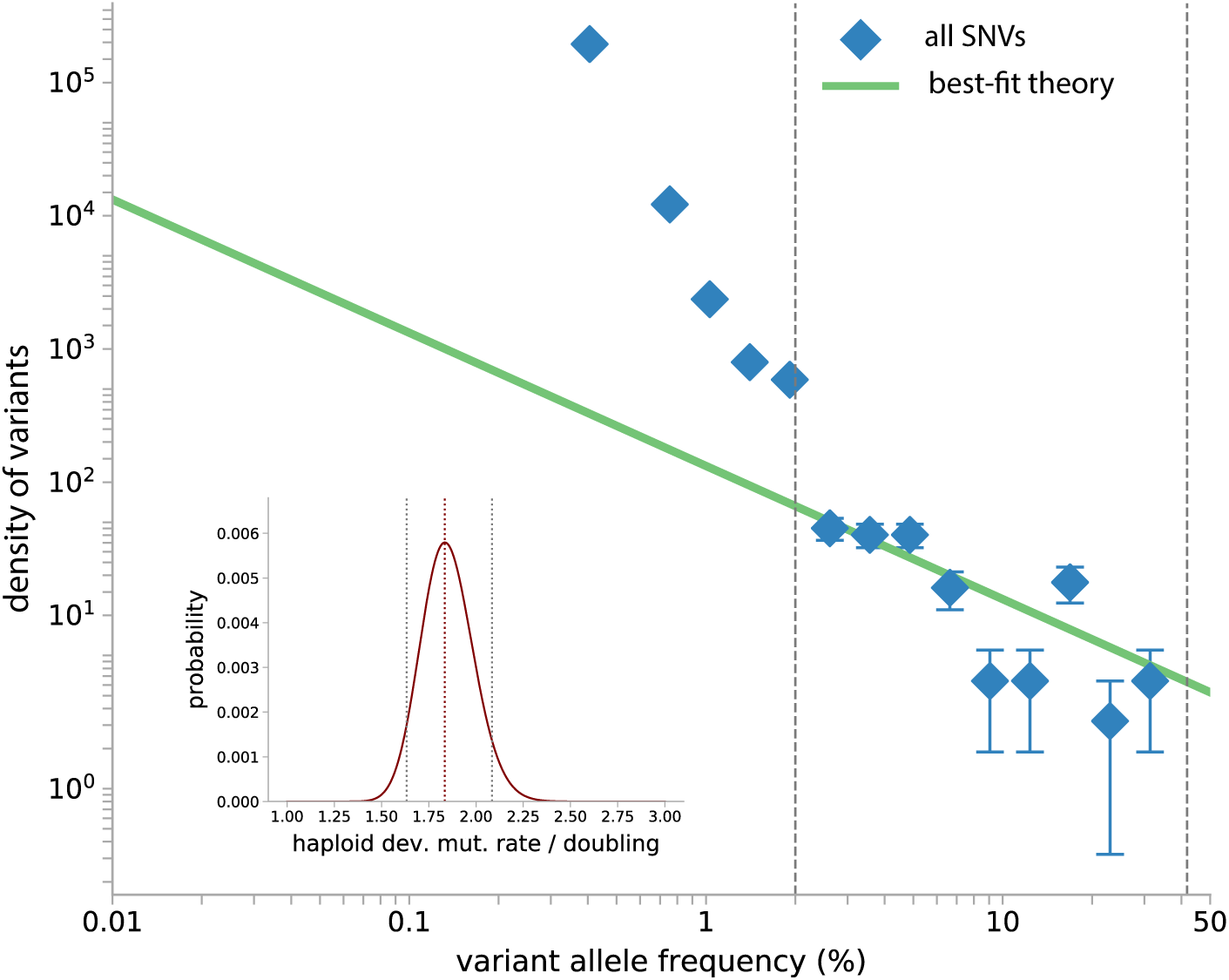
Developmental mutation rates averages to 2 − 4 across entire genome per cell doubling. Best-fitting to the reverse cumulative density (blue diamonds) - not normalized by mutation rates - is performed for SNV data obtained through single-cell whole genome sequencing down to *f*′ = 2% VAF and below ∼ 42% (grey dashed lines). The haploid developmental mutation rate is 1.8 (95% CI interval: 1.6 − 2.1) per cell doubling (green line). Inset: Likelihood plot of best-fit.

### C. Drift-selection boundary and self-consistency of neutral parameters in blood

The drift-selection boundary in healthy blood is evident at VAF 0.2% in healthy blood according to variant data from Razavi et al. ^15^ (Figure S6). Because the vast majority of missense and synonymous variants are dominated by neutral drift below Ψ ≈ 0.2%, i.e. the VAF at which the abundance of mutants driven by drift become less than that of hitchhiking mutants, only the part of the VAF spectrum of both below 0.2% are best-fitted with a three-parameter fit to the neutral drift prediction to obtain *Nτ* = 4.9 × 10^4^ years, haploid synonymous mutation rate 8.1×10^−4^ and haploid missense mutation rate 1.8 × 10^−3^ per year. The inferred value of the synonymous mutation rate is in good agreement with what would be estimated based on trinucleotide context (≈3 × 10^−3^ per year) ^3,17^ considering that false negative rates increases at low VAFs due to detection limits. The inferred combination of parameters *Nτ* is consistent with that found in other studies ^3,17^. It is worth noting that the inferred estimate is within a factor of ∼ 2 of the estimate from Watson et al. ^17^ and both are within the range of estimates in literature ^3^. The inferred missense mutation rate is 2.1 times the inferred synonymous mutation rate, which is in reasonable agreement with the estimated ratio of nonsynonymous and synonymous sites on the sequencing panel scaled from the Illumina Myeloid panel (≈2.6). It was checked that fitting the two VAF spectra separately for *Nτ* yields values (synonymous: *Nτ* ≈5.0 × 10^4^ years, missense: *Nτ* ≈4.8 × 10^4^ years) very close to the three-parameter fit that considers missense and synonymous VAF spectra simultaneously. The consistency between data and with the neutral drift prediction strongly suggests that the system transitions from being dominated by genetic hitchhiking to being dominated by drift below 0.2% VAF where the theory captures the shape of the spectra precisely.

For our analyses on healthy blood, the passenger theory is fitted based on a previous inference of *Nτ* = 1 × 10^5^ years ^17^ for consistency, which is also within the range of literature values ^3^, and only applied to data above the drift limit Ψ. In healthy oesophagus, the drift-selection boundary occurs below detection limits in Martincorena et al. ^5^ according to best-fits (*Nτ* = 1 × 10^4^ years) performed on the TP53 nonsynonymous VAF spectrum (Supplementary note 5B). Therefore, the passenger theory is fitted to healthy oesophagus data above the drift limit which, in this case, does not exclude any data from the original study after trimming (Supplementary note 5D). Because data in Razavi et al. ^15^ is normalized based on a genomic region size of 2.13 Mb, quoted in the original paper ^15^, which is an upper bound to the size of the coding region, inferred mutation rates and *Nτ* estimates are lower bounds from this analysis.

**Fig. S6.**
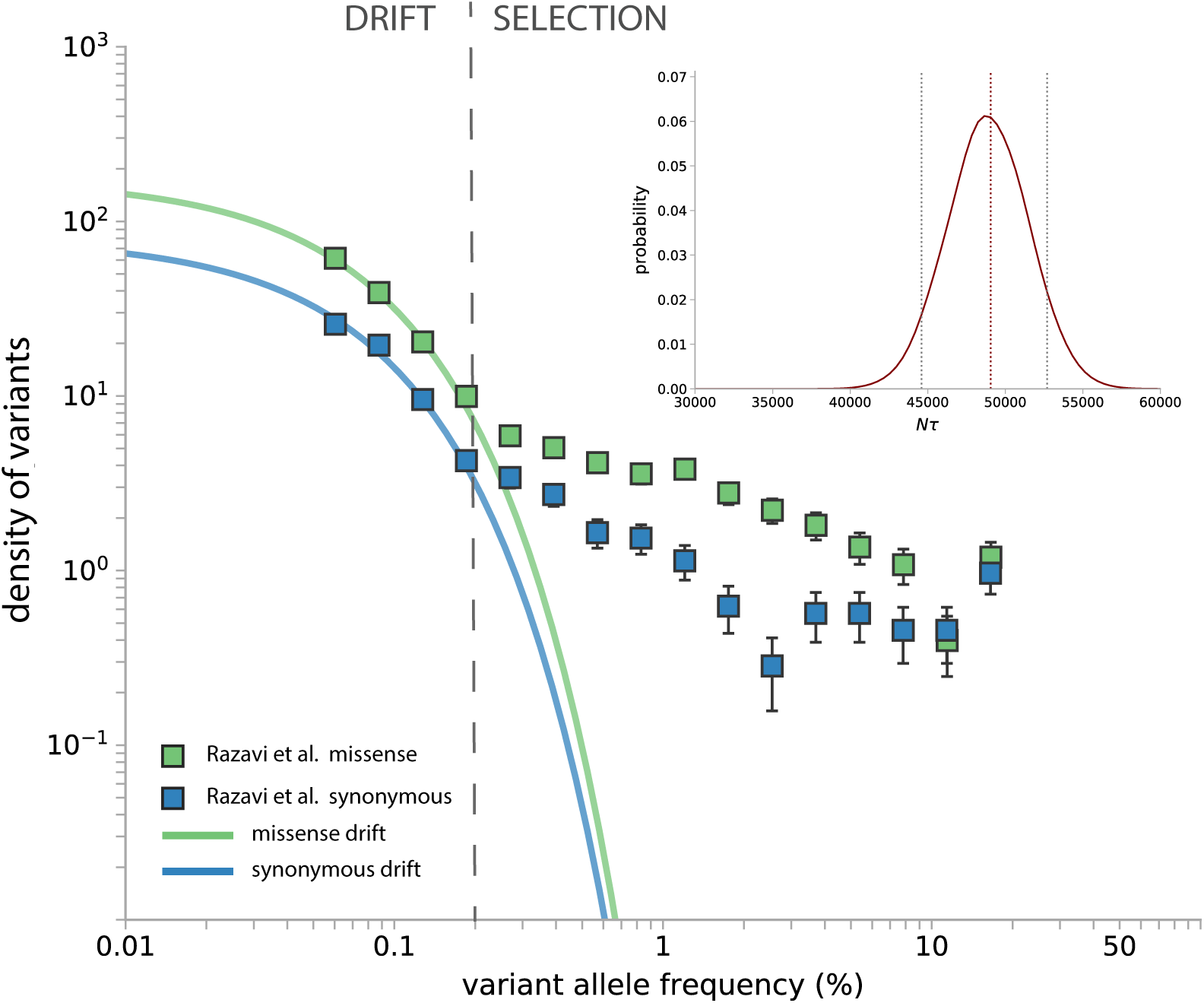
Drift-selection boundary is evident in Razavi et al. data. Variant data of all synonymous (blue squares) and missense variants (green squares) between VAF 0.05% and 20% from Razavi et al. ^15^ is plotted without normalizing by study-specific mutation rates. Transition from drift to selection is evident at VAF ≈ 0.2%. Best-fit predictions are shown (*Nτ* = 4.9 × 10^4^ (95% CI= 4.5 × 10^4^ − 5.3 × 10^4^) years, haploid synonymous mutation rate = 8.1 × 10^−4^ and haploid missense mutation rate = 1.8 × 10^−3^ per year).

### D. Predicting passengers based on the top 20 quantified high-fitness ‘drivers’

In order to assess how sensitive our predictions of the number of synonymous passengers are to the DFE, here we quantify what distribution of synonymous passenger mutations would be expected if hitchhiking occurred alongside 20 of the fittest variants in clonal haematopoiesis, whose distribution of fitness effects can be inferred directly from data whereas ‘driver’ mutations of moderate fitness cannot as they are detected less frequently. The fitness effects and mutation rates of 20 common high-fitness variants were quantified by analysing their VAF distributions in large cohorts of individuals in previous work ^17^. The estimates for fitness effect and mutation rate of these 20 variants form a distribution of mutation rates over fitness effects which does not require assuming a particular functional form for the DFE (Figure S7A). This empirical distribution can be used to predict the expected density of synonymous passengers using Equation 12 by assuming summation instead of integration over the fitness distribution. As expected, the predicted passenger VAF spectrum underestimates observed synonymous variant densities (Figure S7B). This observation that the common high fitness ‘drivers’ cannot account for most hitchhiking is further evidenced by the correlation of VAFs for the largest synonymous variant and the largest nonsynonymous variant shown in Figure 2C, where it is clear that a reasonable proportion of the synonymous variants at VAFs >10% do not have a putative ‘driver’ anywhere in the panel, let alone in one of the common 20 driver positions.

**Fig. S7.**
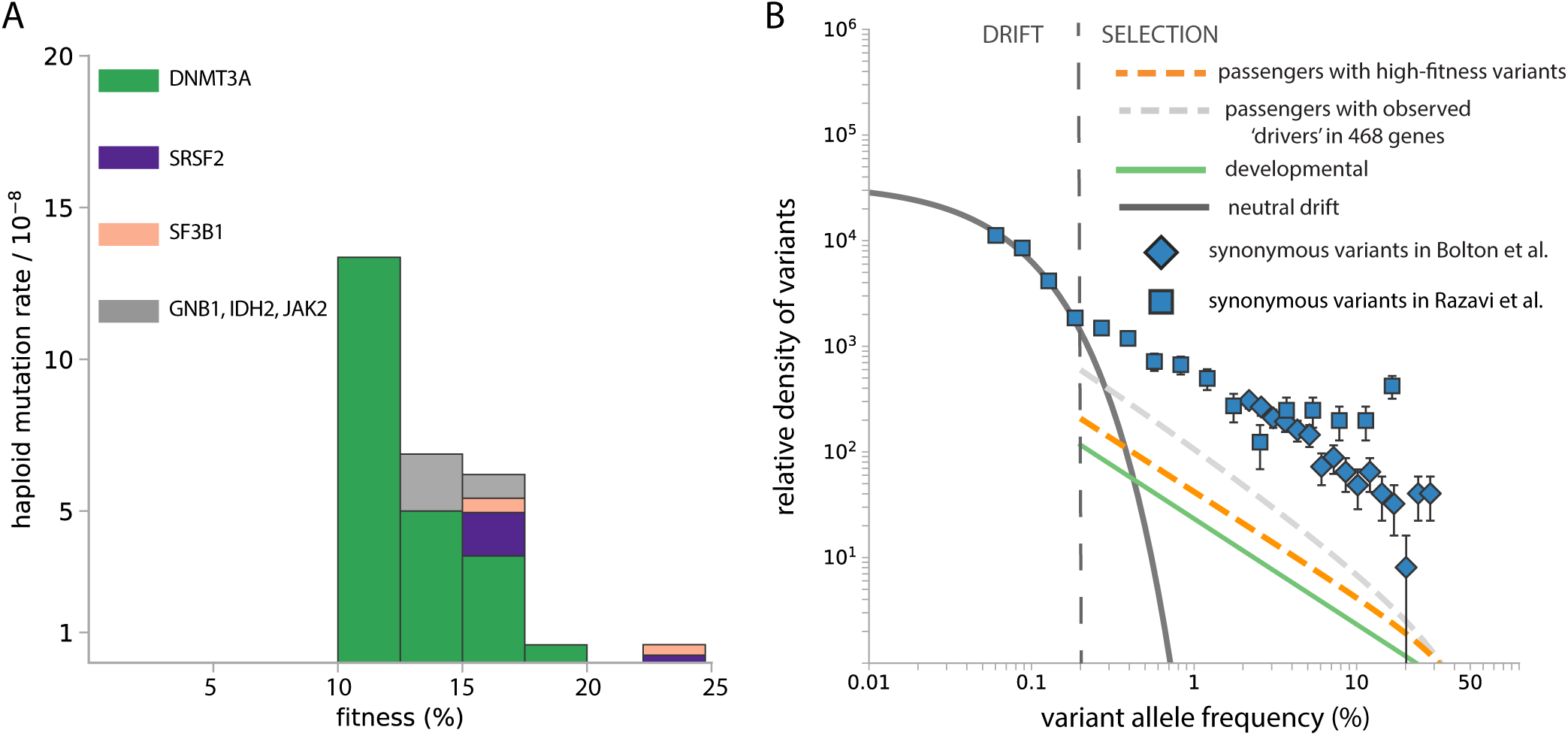
Predicting passengers VAF spectra due to 20 common high-fitness variants in clonal haematopoiesis ^17^. (A) The fitness landscape of the top 20 discrete drivers in ^17^ binned in 2.5% fitness intervals. These top variants belong to DNMT3A, SRSF2, SF3B1, GNB1, IDH2, and JAK2. (B) The shape of the predicted passenger VAF spectrum is consistent with 1*/f* ^2^ only considering strong drivers but still underestimates the observed synonymous variant abundance at high and intermediate VAFs.

### E. Fitness landscape model

To make predictions for the observed hitchhiker VAF density, we used the parametrized distribution of fitness effects as described below. Different mutations affect the survival advantages of the cell to different extents and only a small portion of them confer strong survival advantage to the cell. There are many ways of parametrizing the fitness landscape, one of which is the exponential of a power distribution, which captures the strongly decreasing prevalence of mutations with high fitness:

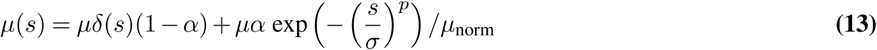

Where 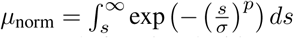. The free parameters are *p, σ* and *α*, the latter representing the fraction of variants that are non-neutral (functional ‘drivers’) among all nonsynonymous variants in the panel (haploid nonsynonymous mutation rate *µ* = 1.6 × 10^−3^ per year across the panel). This DFE captures ‘driver’ with relatively large fitness effects and a large fraction that behave neutrally and do not experience positive selection.

In the Bolton et al. ^14^ cohort, 269 nonsynonymous variants were detected among the 399 individuals sequenced on the latest panel version (covering 468 cancer-associated genes). Only 12 types of variants were recurrent (i.e. detected more than once) and therefore presents sufficient data for inferring fitness effects for each specific variant based on occurrence times or VAF spectra. The distribution of fitness effects (DFE) of all 269 nonsynonymous variants is assumed to consist of a peak at *s* = 0 and a general functional form as above (Equation 13), where *p >* 1, *σ* and *α* are free parameters.

**Fig. S8.**
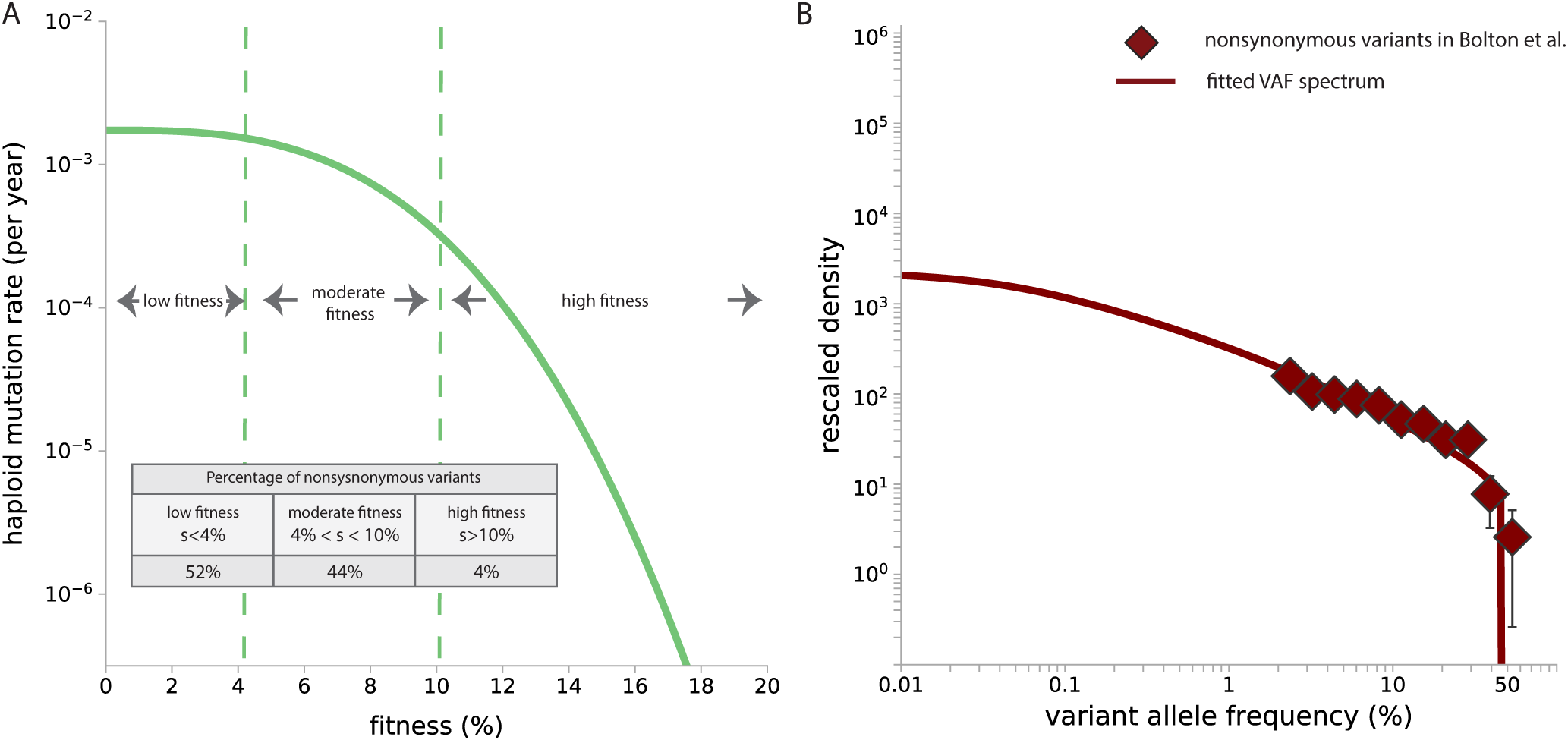
Distribution of fitness effects of nonysynonymous variants in Bolton et al. ^14^. Only individuals sequenced on the 468-gene panel are considered in the fitting of the DFE. (A) Best-fit DFE (*p* = 2.9, *σ* = 0.08, *α* = 0.02) implies 2% of all nonsynonymous variants on the panel are functional (functional driver mutation rate is 3.5 × 10^−5^ per year per haploid - the exact value is dependent on the shape of DFE), among which 4% are of high fitness (*s >* 10%), 44% are of moderate fitness (4% *< s <* 10%) and 52% are of low fitness (*s <* 4%). (B) The best-fit nonsynonymous VAF spectrum predicted for the 269 nonsynonymous variants based on the exact distribution of ages.

### F. Contribution to the passenger prediction from different parts of the DFE and different ages

Based on the form of DFE fitted from all nonsynonymous variants in section E, the passenger VAF spectrum of the Bolton et al. ^14^ cohort can be predicted based on their ages. The median age of the cohort is 67.5 years (lower quartile: 61.5 years; upper quartile: 74.9 years). Different parts of the DFE contribute to the final passenger VAF prediction to different extents depending on the person’s age (Figure S9). For the age distribution in the Bolton cohort, the fraction of driver mutations with fitness between 9% and 15% is the most dominant at VAFs above 2%. At these fitnesses, the VAF prediction shows a characteristic scaling of 1*/f* ^2^. The overall passenger prediction has a similar scaling but falls off slightly faster due to contributions from remaining parts of the DFE.

**Fig. S9.**
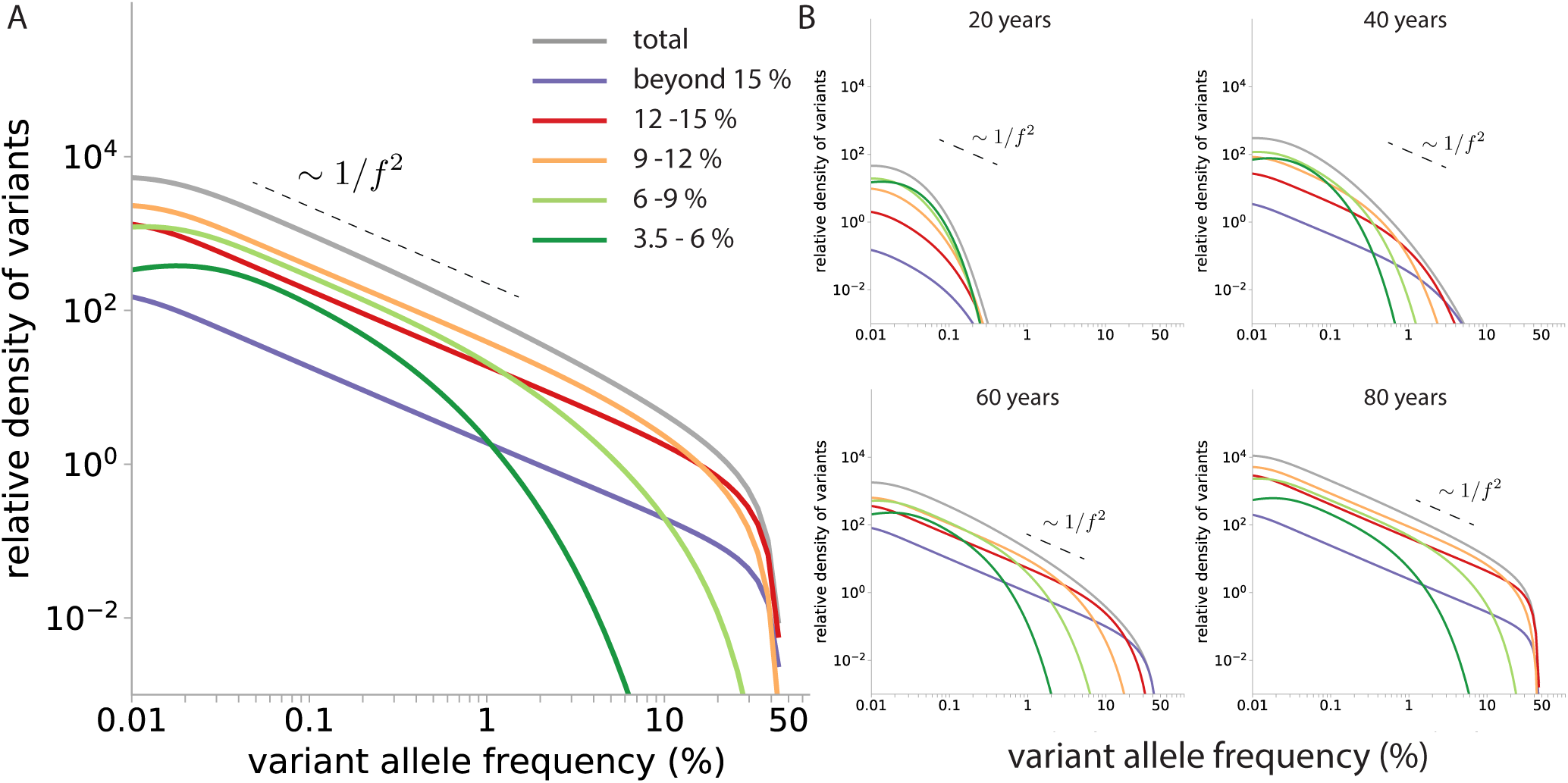
Predicting the passenger VAF spectra contributed by different parts of the DFE. (A) For the age distribution of the 590 individuals in Bolton et al. ^14^, the passenger VAF prediction above VAF *>* 2% is dominated by the fraction of ‘driver’ mutation with fitness 9% *< s <* 15% for the inferred DFE in healthy blood (Supplementary note 3E). (B) The contribution of ‘driver’ mutations with different fitness in the DFE changes for each person as they age. The total (grey line) represents the total passenger VAF spectrum contributed by ‘driver’ mutation fitness *s >* 3.5%.

### G. Inferring total ‘driver’ mutation rate and DFE in healthy blood

Similar to our stochastic simulation validations, a one-parameter fit is performed to the reverse cumulative of the synonymous VAF spectrum above the drift limit Ψ = 2 × 10^−3^, i.e. the VAF at which the abundance of mutants driven by drift become less than that of hitchhiking mutants (section C), and within the VAF window 2 − 35%, beyond which the detectability of variants is limited by sequencing depth and cohort size. Fitting is performed using the ‘Nelder-Mead’ algorithm across bins in log-scale with respect to reverse cumulative densities and weighted by sampling error. Despite the inferred haematopoietic stem cell (HSC) population size based on the observed drift-selection boundary (section C, *Nτ* = 4.9 × 10^4^ years), the parameter is fixed to be the slightly larger value *Nτ* ≈ 1 × 10^5^ years inferred in previous work for consistency ^17^. Study-specific mutation rate and fraction of synonymous sites are scaled from our previous work ^17^ according to the panel size of the study.

We found that the passenger prediction based on the 468 cancer-associated genes covered by the largest panel used in Bolton et al. ^14^, which are not restricted to being related to clonal haematopoiesis only, underestimates the numbers of high-frequency synonymous variants detected in healthy blood (Figure S10, orange dashed line). The total ‘driver’ haploid mutation rate that drives all clonal expansions is 3.4 × 10^−4^ (95% CI: 3.0 × 10^−4^ −3.8 × 10^−4^) per year (Figure S10, orange solid line) for the DFE form used (Figure S8), implying that there are ∼ 9 fold more unobserved ‘drivers’ compared to the contribution from the cancer-associated genes alone. This fold estimate of unobserved ‘drivers’ is only a lower bound as the Bolton et al. ^14^ cohort is sequenced on three different panels. We used the largest panel that encompasses all genomics regions covered in the two previous panel versions for normalization (the smallest panel is ∼ 80% of the largest panel in terms of base pairs). As the result, the actual synonymous VAF densities would be higher than what is considered here and would correspond to a higher fold of unobserved ‘drivers’.

**Fig. S10.**
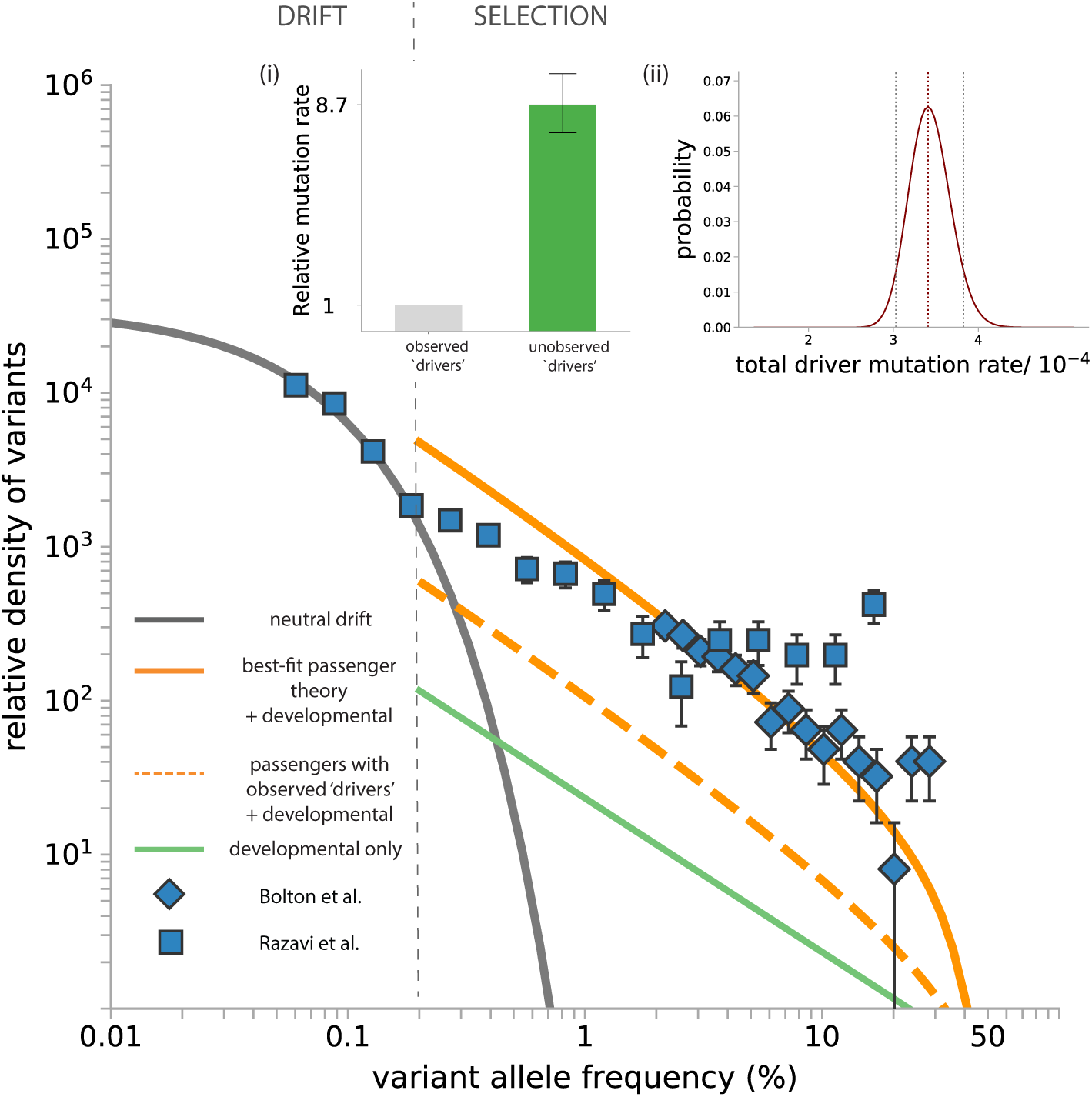
Synonymous variants in healthy blood. Synonymous variant data from ^14,15^ are compared to the passenger prediction for those that hitchhiked with observed ‘drivers’. The prediction for passengers based on 468 cancer-associated genes (orange dashed line, passengers with observed ‘drivers’) underestimates numbers of high-frequency synonymous variants in healthy blood. Developmental mutations are estimated to be a small contribution (green line). The best fit (orange solid line, best-fit passenger theory + developmental) total driver haploid mutation rate per year for the DFE form used (Figure S8) is 3.4 × 10^−4^ (95% CI: 3.0 × 10^−4^ − 3.8 × 10^−4^) from fitting to Bolton data alone. The neutral drift prediction is based on estimates of HSC population size and study-specific synonymous mutation rate inferred from the drift-selection boundary (section C). The predictions are overlaid with data from Bolton et al. ^14^ as well as Razavi et al. ^15^. (i) There are 8.7 (95% CI: 7.6 − 9.9) fold more unobserved ‘drivers’ compared to the contribution from the 468 cancer-associated genes. (ii) Likelihood plot shows the 95% confidence interval for one-parameter fitting of the total functional ‘driver’ mutation rate across the fitness landscape with respect to the DFE form in section E.

### E. Sensitivity to functional form of the DFE

It is natural to question whether and to what extent the specific form of fitness landscape (DFE) chosen for parametrization (Supplementary note 3E) affects our predictions and if constraining the shape of the DFE impacts our analysis. The exact same analysis in Supplementary note 3G is carried out by assuming the DFE form according to Equation 13 but for fixed *p* = 2, 3, 4 (Figure S11, S12 and S13), where *σ* and *α* remain free parameters.

It is observed that the precise form of the DFE does not change the quantitative predictions that there are 8 − 9 fold more unobserved ‘drivers’ compared to contribution from 468 cancer-associated genes though it does have a (relatively modest) quantitative effect on the exact numbers. As expected, the inferred total functional ‘driver’ mutation rate varies for different specific DFE forms as it represents the rate of possible ‘driver’ mutation summed across all fitness effects and is therefore influenced by the ranges of the DFE (predominantly small *s*) that do not have large contributions to the passenger spectrum or the nonsynonymous ‘driver’ mutation spectrum. However, the inferred fold of unobserved ‘drivers’ is relatively constant across different DFE forms.

**Fig. S11.**
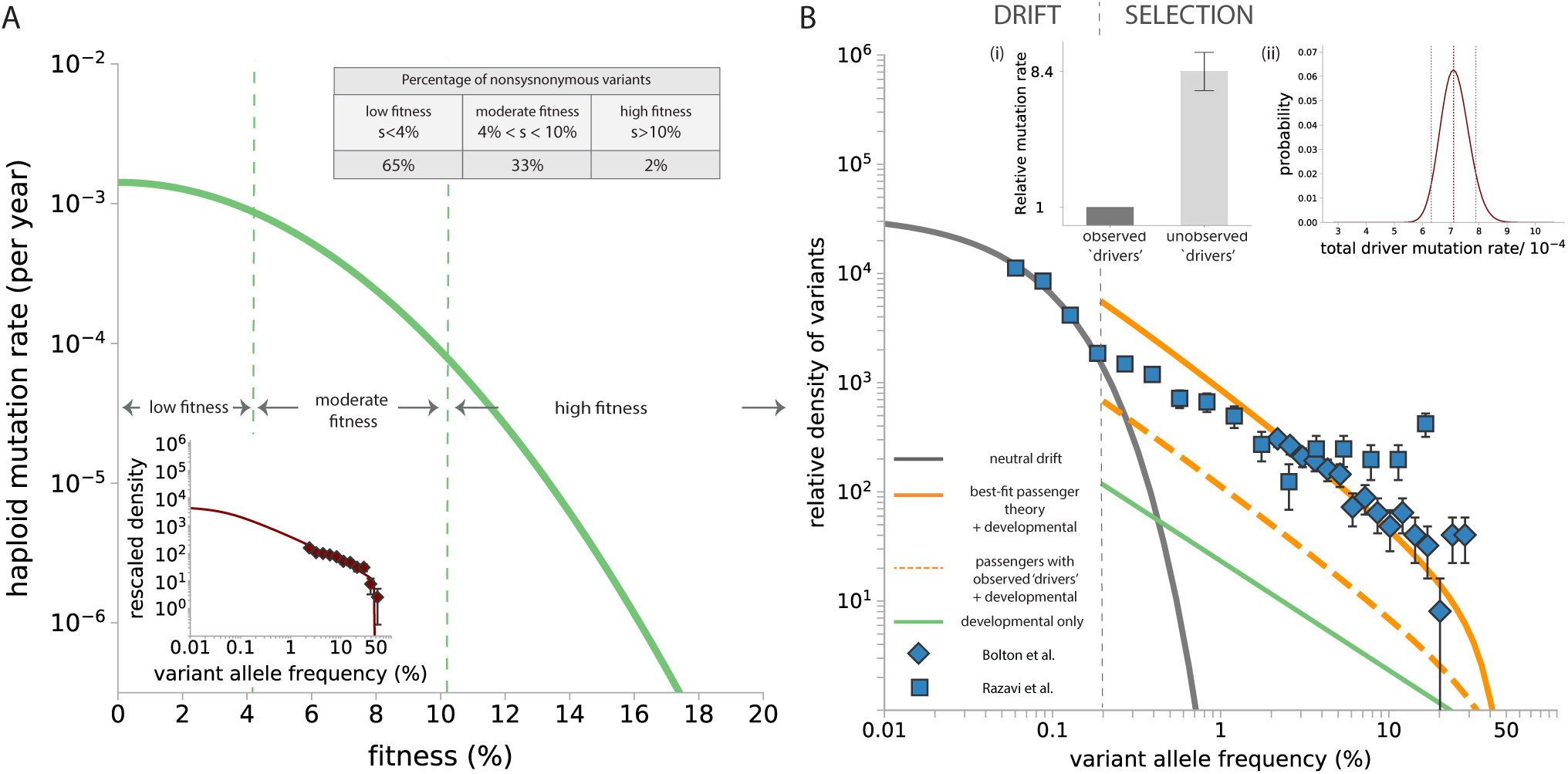
DFE form with *p* = 2. (A) Parameters of the DFE (*p* = 2, *σ* = 0.06, *α* = 0.05) are estimated from the nonsynonymous VAF spectrum in ‘Bolton et al.’ ^14^. Among variants that are non-neutral (which constitute a haploid functional ‘driver’ mutation rate 7.6 × 10^−5^ per year), 2% are of high fitness (*s >* 10%), 33% are of moderate fitness (4% *< s <* 10%) and 65% are of low fitness (*s <* 4%). Inset: The best-fit nonsynonymous VAF spectrum predicted for the 269 nonsynonymous variants based on this DFE. (B) Theoretical predictions for neutral drift (grey line), passengers (orange lines) and developmental mutations (green line) are compared with observed data in ‘Bolton et al.’ ^14^ (blue diamonds) and “Razavi et al.’ ^15^ (blue squares). (i) There are 8.4 (95% CI: 7.3 − 9.4) fold more unobserved ‘drivers’ compared to the contribution from the 468 cancer-associated genes. (ii) Likelihood plot shows the 95% confidence interval for one-parameter fitting of the total functional ‘driver’ mutation rate across the fitness landscape with respect to this DFE form where *p* = 2.The best-fit total functional ‘driver’ mutation rate is 7.1 × 10^−4^ (95% CI: 6.3 × 10^−4^ − 7.9 × 10^−4^) per year (haploid).

**Fig. S12.**
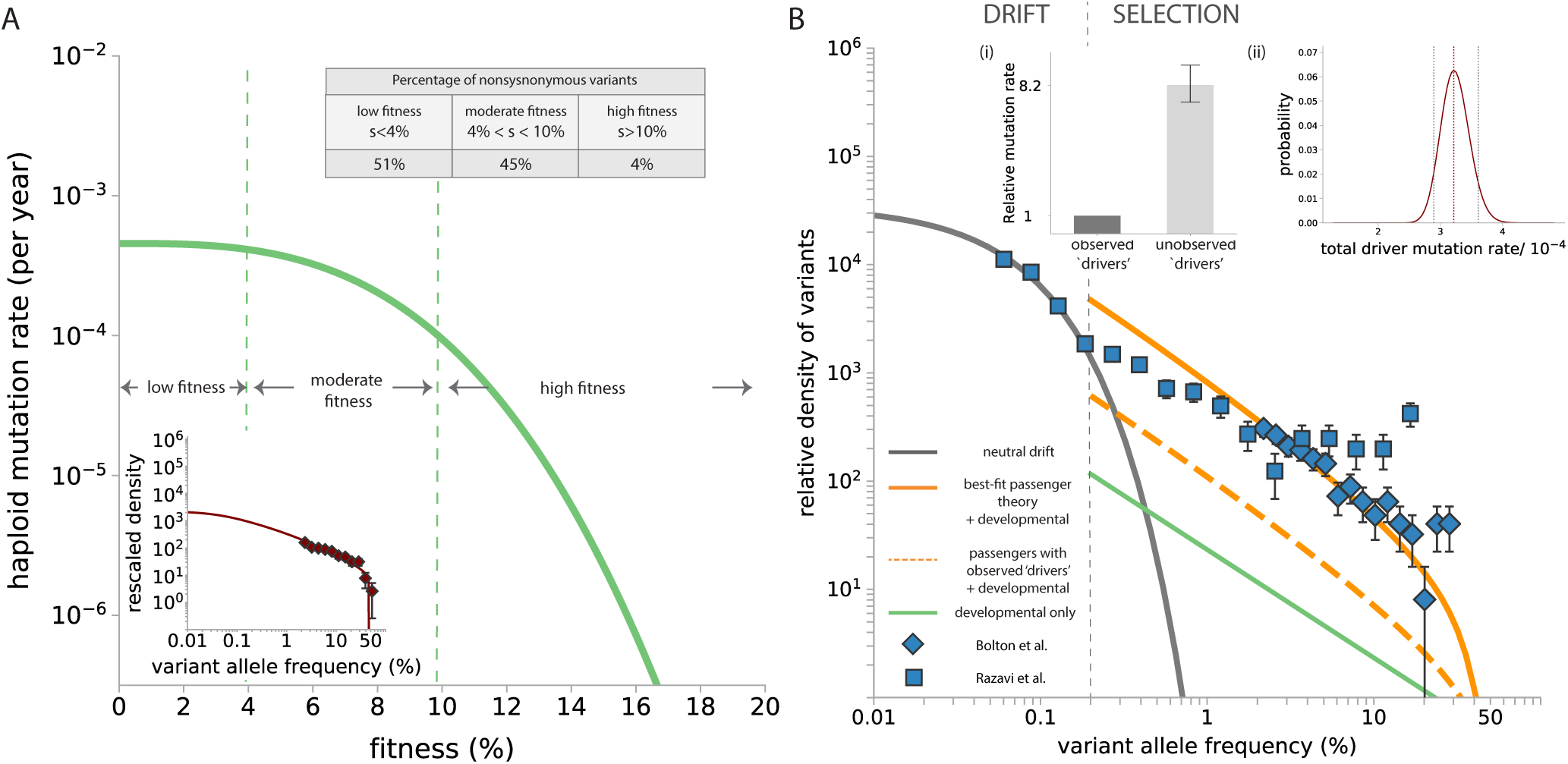
DFE form with *p* = 3. (A) Parameters of the DFE (*p* = 3, *σ* = 0.086, *α* = 0.02) are estimated from the nonsynonymous VAF spectrum in ‘Bolton et al.’ ^14^. Among variants that are non-neutral (which constitute a haploid functional ‘driver’ mutation rate 3.5 × 10^−5^ per year), 4% are of high fitness (*s >* 10%), 45% are of moderate fitness (4% *< s <* 10%) and 51% are of low fitness (*s <* 4%). Inset: The best-fit nonsynonymous VAF spectrum predicted for the 269 nonsynonymous variants based on this DFE. (B) Theoretical predictions for neutral drift (grey line), passengers (orange lines) and developmental mutations (green line) are compared with observed data in ‘Bolton et al.’ ^14^ (blue diamonds) and “Razavi et al.’ ^15^ (blue squares). (i) There are 8.2 (95% CI: 7.3 − 9.3) fold more unobserved ‘drivers’ compared to the contribution from the 468 cancer-associated genes. (ii) Likelihood plot shows the 95% confidence interval for one-parameter fitting of the total functional ‘driver’ mutation rate across the fitness landscape with respect to this DFE form where *p* = 3.The best-fit total functional ‘driver’ mutation rate is 3.2 × 10^−4^ (95% CI: 2.9 × 10^−4^ − 3.6 × 10^−4^) per year (haploid).

**Fig. S13.**
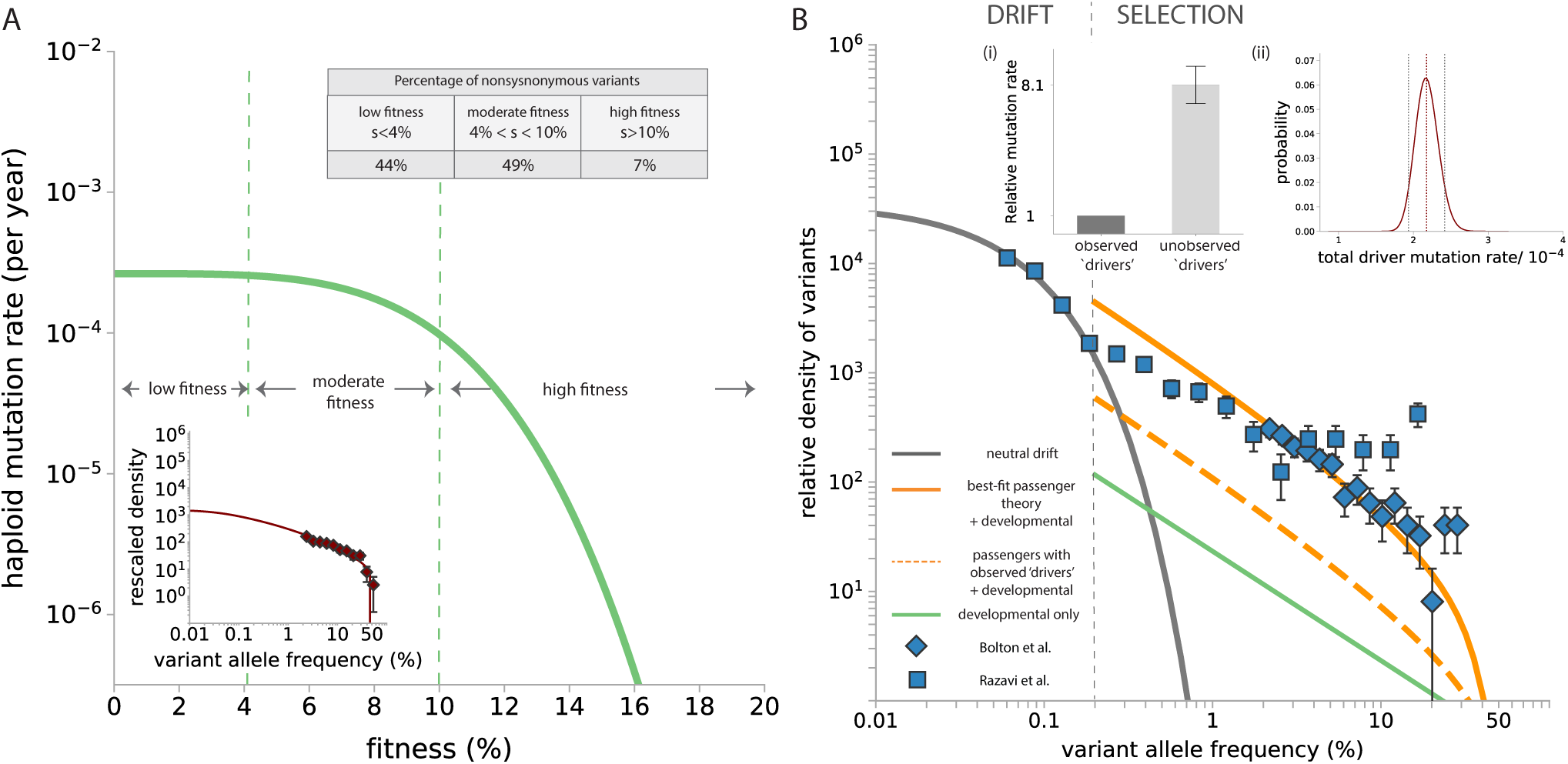
DFE form with *p* = 4. (A) Parameters of the DFE (*p* = 4, *σ* = 0.10, *α* = 0.015) are estimated from the nonsynonymous VAF spectrum in ‘Bolton et al.’ ^14^. Among variants that are non-neutral (which constitute a haploid functional ‘driver’ mutation rate 2.4 × 10^−5^ per year), 7% are of high fitness (*s >* 10%), 49% are of moderate fitness (4% *< s <* 10%) and 44% are of low fitness (*s <* 4%). Inset: The best-fit nonsynonymous VAF spectrum predicted for the 269 nonsynonymous variants based on this DFE. (B) Theoretical predictions for neutral drift (grey line), passengers (orange lines) and developmental mutations (green line) are compared with observed data in ‘Bolton et al.’ ^14^ (blue diamonds) and “Razavi et al.’ ^15^e (blue squares). (i) There are 8.1 (95% CI: 7.0 − 9.0) fold more unobserved ‘drivers’ compared to the contribution from the 468 cancer-associated genes. (ii) Likelihood plot shows the 95% confidence interval for one-parameter fitting of the total functional ‘driver’ mutation rate across the fitness landscape with respect to this DFE form where *p* = 4.The best-fit total functional ‘driver’ mutation rate is 2.2 × 10^−4^ (95% CI: 1.9 × 10^−4^ − 2.4 × 10^−4^) per year (haploid).

### I. Correlation between largest synonymous and nonsynonymous mutant size

If the highest VAF ‘driver’ mutation is found at a higher frequency than the highest VAF synonymous variant in an individual (Figure 2C, shaded region), the ‘driver’ mutation observed is likely causing the hitchhiking (Figure S14A). In contrast, if the highest nonsynonymous VAF is lower than the highest synonymous VAF within an individual, the putative ‘driver’ mutations most likely could not have caused the hitchhiking event unless hitchhiking happened via the ‘neutral-first’ route.

In Bolton et al. ^14^ 73% of individuals harbouring synonymous variants lie below the diagonal (Supplementary Figure S14B, unshaded region), implying that the putative ‘driver’ (nonsynonymous mutations in the 468 cancer-associated genes) is not accountable for the clonal expansion that drove the highest VAF passenger, i.e. large fractions of major clonal expansions are driven by ‘drivers’ outside of these genes. To support this we also ran simulations according to the age distribution in Bolton et al. ^14^, where the driver mutation rates and neutral mutation rates are the same as the haploid rates of the 468-gene panel version, assuming the ratio of observed versus unobserved ‘drivers’ is 1 to 5 and recorded only mutations above the detection limit in Bolton et al., i.e. *>* 2% VAF (Figure S14A). Simulations show 68% individuals lie below the diagonal which is in good agreement with data.

**Fig. S14.**
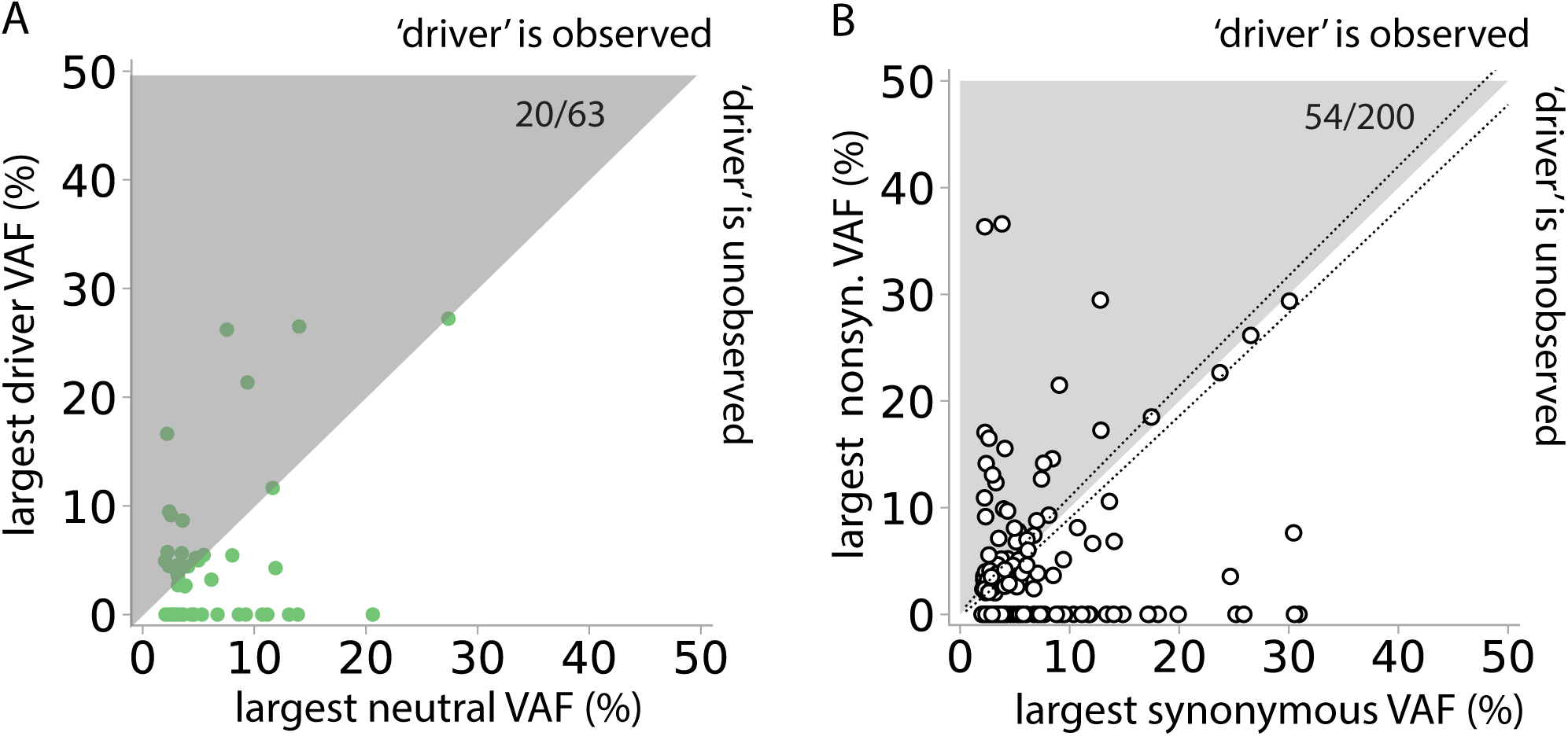
VAF correlation in healthy blood. (A) Green points represent simulated results (*τ* = 1 year). The highest VAF neutral and driver mutations were recorded in each simulation run according to the DFE inferred from blood (Supplementary note 3G) for 590 individuals (ran twice) with the exact same age distribution as the Bolton et al. ^14^ cohort, and neutral mutation rate is equal to the study-specific haploid synonymous mutation rate in Bolton et al. Only mutations above detection limit, i.e. 2% VAF and one in every six drivers was recorded in each individual (i.e. unobserved ‘drivers’ are 5 folds of observed ‘drivers’). Hence only 63 individuals resulted in detectable neutral mutations. The largest observed ‘driver’ and the largest neutral mutation VAF are plotted and 32% (20*/*63) lie above the diagonal. (B) In Bolton et al. ^14^, there are 200 untreated patients who harbour at least 1 synonymous variant before data trimming. 54 harbour a nonsynonymous variant in any gene on the MSK-IMPACT panel that is at a higher VAF than their highest VAF synonymous variant. Most data points are expected to lie below the diagonal if most synonymous variants hitchhike with variants outside these genes, which is indeed the case as only 54 data points are above the diagonal (54*/*200 = 27%). The diagonal dashed lines indicate the upper and lower error due to sampling noise for exactly same VAFs. Taking into account sampling errors due to finite coverage ∼500x, this fraction ranges from 42 − 67 out of 200 (21 − 33%). This suggests that a significant portion of major clonal expansions are driven by ‘drivers’ outside the 468-gene MASK-IMPACT panel or are caused by more complex mutations.

## Supplementary Note 4: Age dependence of variant allele frequency spectra in data

Age dependence in the synonymous VAF spectra, more apparent in the healthy oesophagus than in healthy blood, indicates somatic contribution to synonymous mutation load at high VAF (Figure S15). Disagreement in quantitative predictions suggest that clonal dynamics are more complex than our model basis. It is possible that developmental mutation rates may be higher, however it alone is unlikely to be sufficient to explain the discrepancy fully. Age dependence of the oesophagus VAF spectra is closer to predictions than the blood VAF spectra in Bolton et al. ^14^ This may be due to having a cohort of individuals that are diagnosed with non-haematological cancers which elevates mutation burden in young individuals in the latter ^14^.

**Fig. S15.**
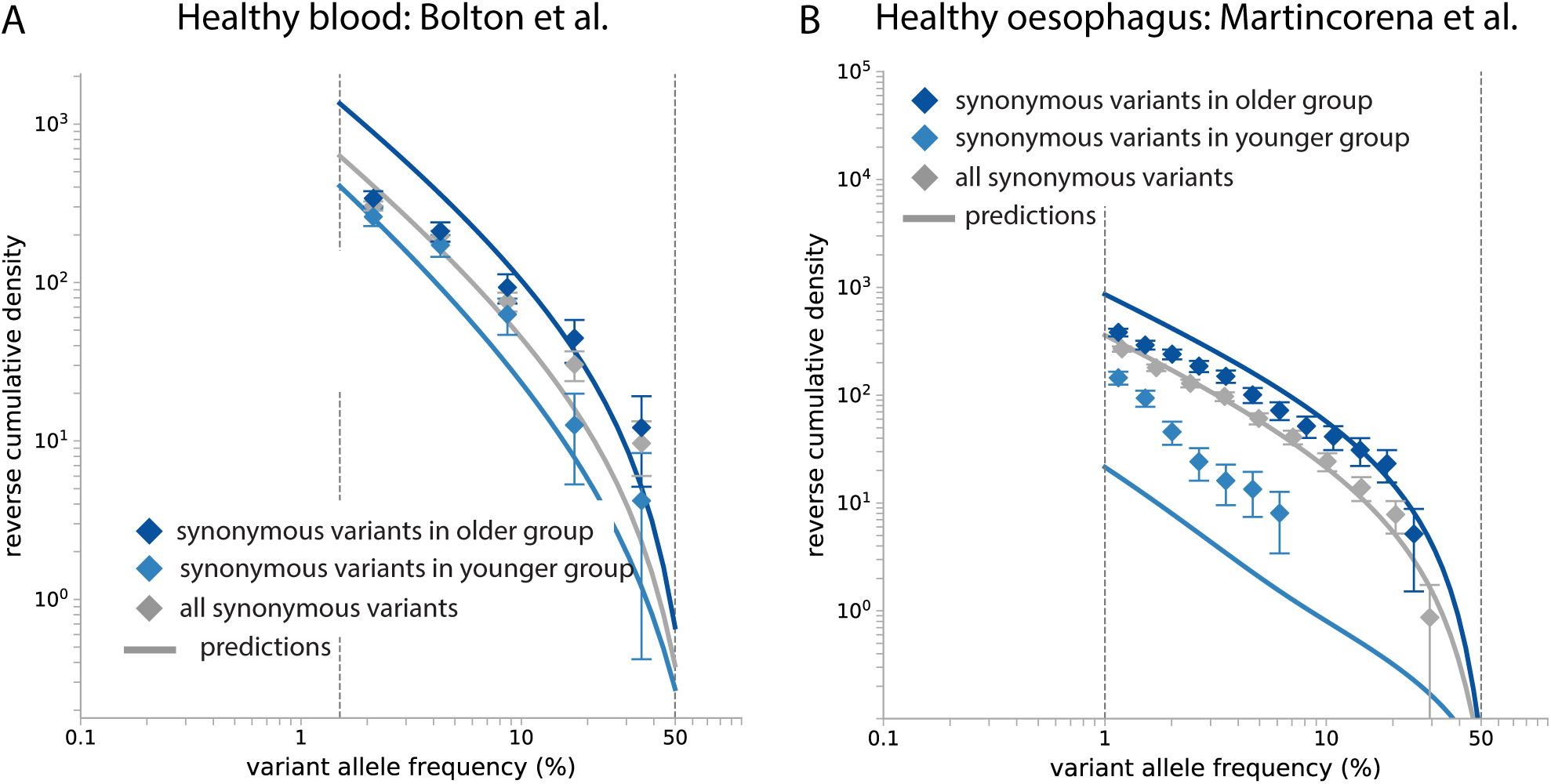
Age dependence of synonymous VAF spectra. Reverse cumulatives of the synonymous VAF spectra are compared with predictions based on the best-fit in our analyses (Supplementary Figure S10 and S17) within the VAF window where we consider there to be low false negative rates (region between the vertical grey lines). The data shows age dependence in both Bolton et al. ^14^ and Martincorena et al. ^5^. (A) Bolton et al. ^14^ is chosen to represent healthy blood because it comprises the vast majority of the entire cohort analyzed in the main text (590 people). There are 194 controls with age 64 − 72 (younger group) and 201 controls with age larger than 72 (older group). (B) The younger age group in Martincorena et al. ^5^ consists of 3 healthy donors of age 21.5, 25.5, 37.5 (sample number is 273) and the older age group consists of 3 oldest healthy donors of age 57.5, 69.5, 73.5 (sample number is 284).

## Supplementary Note 5: Synonymous and nonsynonymous variants in healthy oesophagus

A single cell population maintains squamous epithelial homeostasis in the oesophagus in the absence of slow-cycling stem cells ^47, 48^. This cell population is believed to reside in basal layers and maintains homeostasis by producing on average a daughter cell that replenishes the population and a daughter cell that continues to differentiate, thereby exiting the cell-cycle and moving up supra-basal layers. Our analysis concerns the evolutionary dynamics that happens within such a population.

### A. Data processing

Variant data from post-mortem oesophageal squamous epithelium biopsy samples of nine donors ^5^ were analyzed individually. In each sample, 1 × *Nτ* was first estimated to be 10^4^ years based on the assumption that all synonymous variants observed (blue diamonds, Figure 3) above 1% VAF are passengers (orange dashed curve, Figure 3) and not dominated by drift (grey curve, Figure 3) and later properly fitted using nonsynonymous variants in TP53. Data was trimmed below 1% VAF and above 50% VAF and VAF densities normalized by the number of individuals and the study-specific synonymous mutation rate 2*µ*_syn_ (the panel comprises 74 genes and spans 330 kb). We estimate that around 35 mutations arise per year across the entire genome ^5^. Hence the synonymous mutation rates in oesophagus is estimated at *µ*_syn_ = 6.8 × 10^−4^ per year per haploid for the panel.

We analysed all variant calls after trimming, including collapsed variant calls spanning multiple biopsy samples. Within the same biopsy, mutations shared between samples closer than 10 mm were collapsed in the original study. When more than one biopsy of tissue was available from the same donor, mutations between these biopsies were not merged as distances between biopsies were not available ^5^. Including all collapsed calls in our analysis causes a potential inflation in VAF densities relative to theoretical predictions because we included lineages possibly arising outside of the sample. This is to some extent balanced by the bias introduced by the actual physical clone sizes of these mutants being underestimated. After merging, more than 30% of NOTCH1 (415*/*1251) and TP53 (145*/*451) nonsynonymous calls are mutations detected in multiple biopsy samples in a person and have undergone merging, whereas only a small portion (57*/*603 *<* 10%) of synonymous variant calls in the merged set have been corrected by merging because VAFs associated with synonymous variants are typically smaller. Hence, our analysis of passengers VAF spectra in the merged set is less limited by finite biopsy sample sizes.

### B. Fitness landscape model

The mutational distribution of fitness effects for observed drivers in our analysis consists of two delta-functions corresponding to TP53 and NOTCH1 with rates equal to *µ*_TP53_ and *µ*_NOTCH1_. To estimate that, firstly, *Nτ* is obtained from a 3-parameter fit (*µ*_TP53_, *s*_TP53_, *Nτ*) from the nonsynonymous VAF spectrum of TP53 and from then on fixed at the best-fit value, i.e. *Nτ* ≈1 × 10^4^ years. The other two best-fit parameters are *µ*_TP53_ = 1.4 × 10^−5^ (95% CI : 1.13 − 1.60 × 10^−5^) per year per haploid and *s* = 0.10 per year (95% CI : 0.096 − 0.107) (CI obtained from 2-parameter fit) per year (Figure S16A). Secondly, we perform a two-parameter fit with the nonsynonymous VAF spectrum of NOTCH1 and found the best-fit parameters to be *µ*_NOTCH1_ = 3.3 × 10^−5^ (95% CI : 3.00 − 3.56 × 10^−5^) per year per haploid and *s* = 0.11 per year (95% CI : 0.107 − 0.112) (CI from 2-parameter fit) (Figure S16B).

**Fig. S16.**
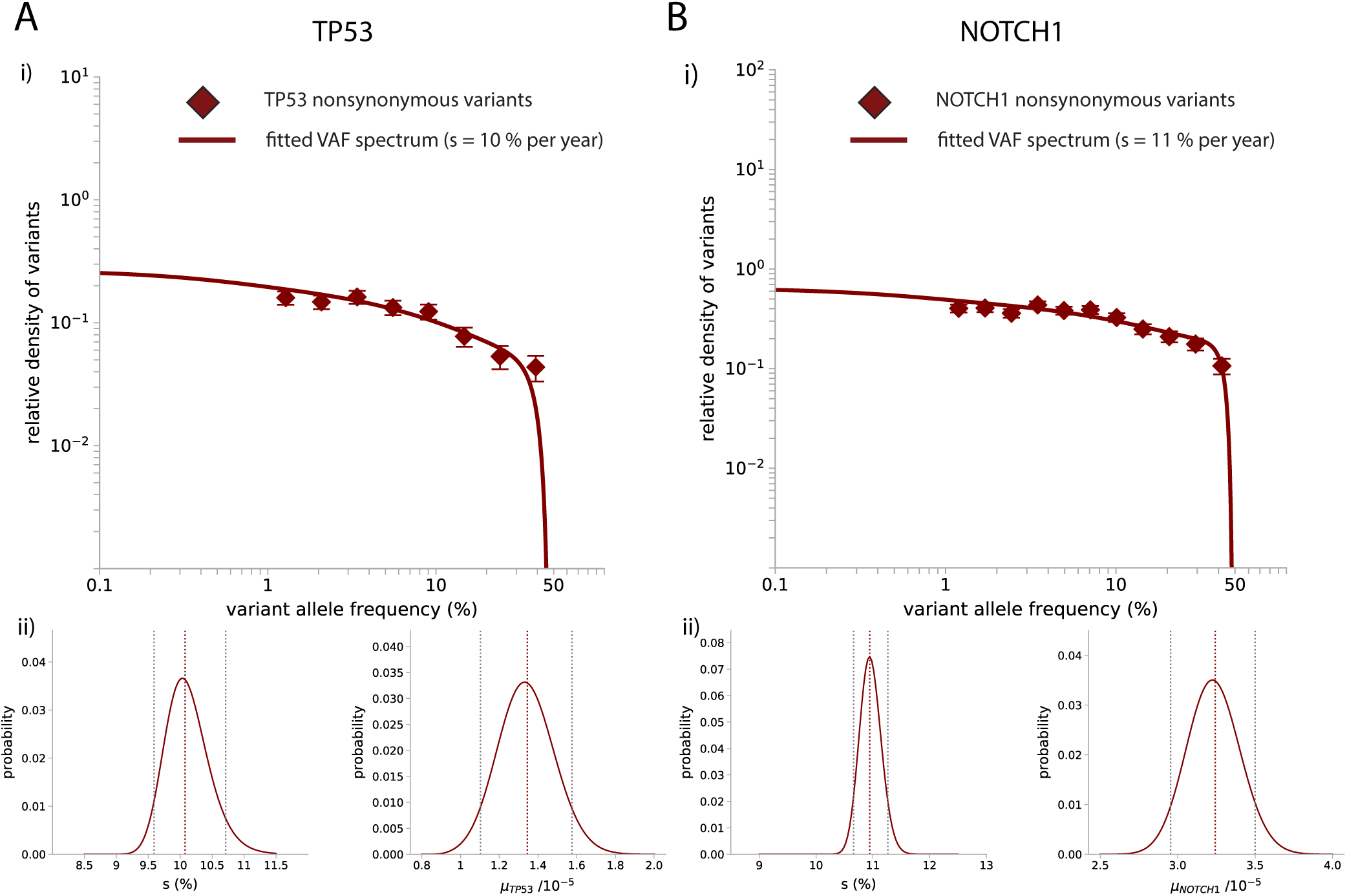
Nonsynonymous variants in healthy oesophagus. (A) Nonsynonymous variants (maroon diamonds) at VAF *>* 1% in TP53 are fitted with predictions for driver mutations (maroon solid line). Based on *Nτ* = 1 × 10^4^ years from a 3-parameter best-fit of the TP53 nonsynonymous VAF spectrum, it is estimated that *µ*_TP53_ = 1.4 × 10^−5^ (95% CI : 1.1 − 1.6 × 10^−5^) per year per haploid and *s* = 0.10 per year (95% CI : 0.096 − 0.107) for TP53. (B) Nonsynonymous variants (maroon diamonds) at VAF *>* 1% in NOTCH1 are fitted with predictions for driver mutations (maroon solid line). Based on *Nτ* = 10^4^ years, it is estimated that *µ*_NOTCH1_ = 3.3 × 10^−5^ (95% CI : 3.0 − 3.6 × 10^−5^) per year per haploid and *s* = 0.11 per year (95% CI : 0.107 − 0.112) (CI from 2-parameter fit) for NOTCH1.

### C. Developmental mutations

Because no robust estimates for developmental mutation rates in healthy oesophagus are readily available, our analysis assumes that early developmental mutations are shared between blood and oesophagus and we applied the same estimate for the developmental mutation rate per bp from Supplementary note 3B. The developmental contribution to the observed synonymous VAF spectrum is also small (Supplementary Figure S17, green line).

### D. Inferring total ‘driver’ mutation rate in healthy oesophagus

The synonymous VAF spectrum was normalized by the study-specific diploid synonymous mutation rate estimate (2*µ*_syn_ where *µ*_syn_ = 6.8 × 10^−4^) after taking into account the fraction of synonymous sites (= 0.354). Assuming all drivers share the same fitness as NOTCH1 and TP53, we fit for the total ‘driver’ mutation rate by considering the reverse cumulative of the spectrum within the VAF window 1 − 40%, beyond which the detectability of variants is limited by sequencing depth or cohort size. Observed synonymous passengers were found to be only 1.7 (95% CI : 1.6 − 1.9) fold more than that contributed by NOTCH1 and TP53, suggesting that NOTCH1 and TP53 account for most of the genetic hitchhiking and only a small remainder is contributed by other genes which are either less frequently mutated or less selected on.

**Fig. S17.**
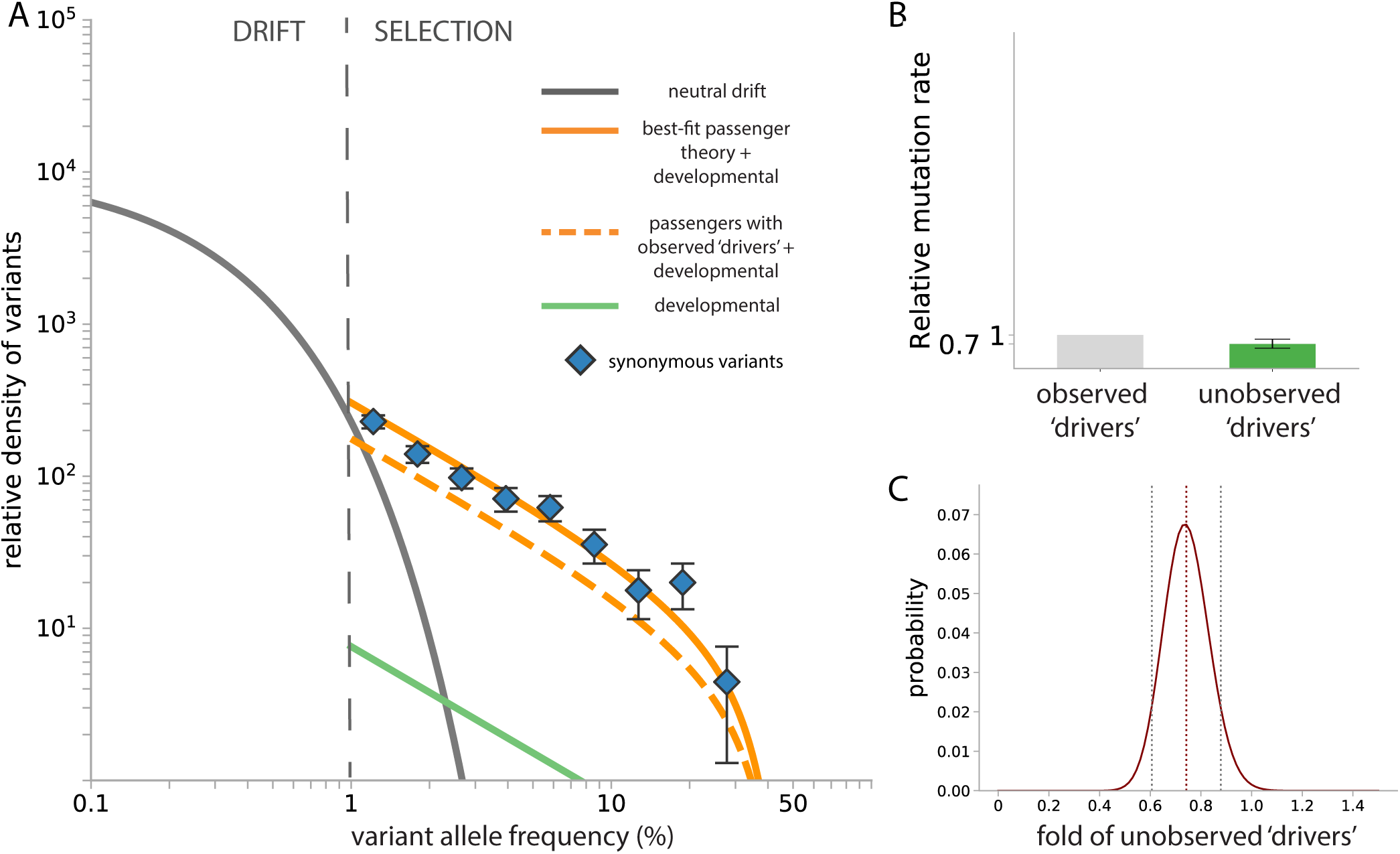
Synonymous variants in healthy oesophagus. (A) Synonymous SNVs reported in ^5^ (blue diamonds) are plotted as VAF densities after normalization by 2*µ*_syn_ where *µ*_syn_ = 6.8 × 10^−4^ is the estimated annual haploid synonymous mutation rate across the sequencing panel. The developmental contribution (green line) is small relative to somatic contributions. The abundance of synonymous passengers predicted driven by NOTCH1 and TP53 ‘drivers’ (orange dashed line) is close to what is observed (blue diamonds). The total genome-wide annual haploid driver mutation rate that fully explains the abundance of passengers (solid orange line) assuming other ‘drivers’ follow the same DFE is found to be 1.7 (95% CI : 1.6 − 1.9) fold more than the contribution from nonsynonymous NOTCH1 and TP53 variants alone. The neutral drift prediction is based on the actual ages of the donor samples and *Nτ* inferred in supplementary note 5B. (B) Relative mutation rates of observed ‘drivers’ and unobserved ‘drivers’ in healthy oesophagus. (C) Likelihood plot for the one-parameter fit for the fold of unobserved ‘drivers’: 0.7 (95% CI : 0.6 − 0.9).

### E. Correlation between largest synonymous and nonsynonymous mutant size

If the highest VAF ‘driver’ mutation is found at a higher frequency than the highest VAF synonymous variant in a sample, the ‘driver’ mutation observed is typically causing the hitchhiking. In contrast, if the highest nonsynonymous VAF is lower than the highest synonymous VAF within a sample, the putative ‘driver’ mutations most likely did not cause the hitchhiking event. In healthy oesophagus, the former scenario is true in 78% of all samples (*n* = 277) with any detected synonymous variants (Supplementary Figure S18), supporting the notion that the putative ‘drivers’ (nonsynonymous mutations in NOTCH1 and TP53) account for most clonal expansions and there is little positive selection outside of these two genes. The observation that over 87% (241*/*277) of samples harbouring synonymous variants also contain either a NOTCH1 or TP53 nonsynonmous SNVs in our trimmed data set, also further supports our conclusion that these two driver genes are responsible for most of the selection in healthy oesophagus.

**Fig. S18.**
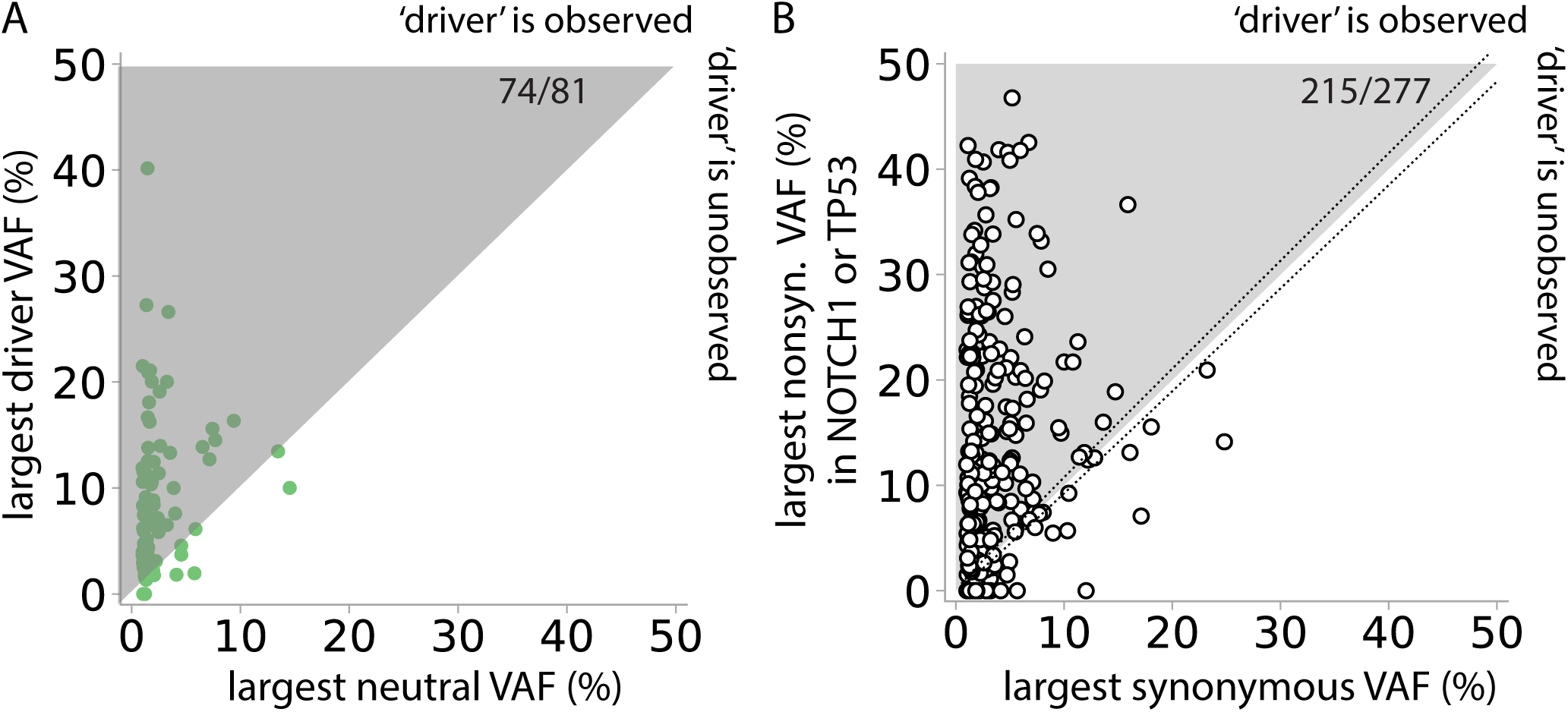
VAF correlation in healthy oesophagus. (A) Green points represent simulated results (*τ* = 1 year). The highest VAF neutral and driver mutations are recorded in each simulation run (*n* = 844, ran twice) with the same age distribution as the Martincorena et al. ^5^ samples. In the simulations mutation rates correspond to the haploid mutation rates of NOTCH1 and TP53 (with corresponding fitnesses) and the study-specific haploid synonymous mutation rate in Martincorena et al. ^5^. The ratio of observed ‘drivers’ versus unobserved ‘drivers’ is set to 1 and only mutations above detection limit, i.e. 1% VAF, are recorded. Simulation results show 81 samples harbour at least one detectable neutral mutation. (B) In Martincorena et al., 277 out of 844 samples contain any detected synonymous variants on the sequencing panel. 215 out of these 277 pairs of data lie above the black diagonal (215*/*277 = 78%). Taking into account sampling errors due to finite coverage ∼ 870x, this fraction ranges from 209 − 222 out of 277 (75 − 80%). The diagonal dashed lines indicate the upper and lower error due to sampling noise for exactly same VAFs.

## Supplementary Note 6: More complex genetic mutations

We considered evidence for selection on the copy number alteration in which the Y chromosome is somatically lost in healthy blood, which occurs in a large fraction of blood in some men ^29^. Analysing data (droplet digital PCR) of Y-loss from ^29^ with our method ^17^ suggests that loss of Y confers a fitness advantage of ∼11.0% (95% CI = 10.7% − 11.3%) per year in the blood and occurs at a rate 2.5 × 10^−6^ (95% CI = 2.1 − 2.9 × 10^−6^) per cell per year. This likely account for part (*<* 10%) of the unobserved ‘drivers’ in our healthy blood analysis.

**Fig. S19.**
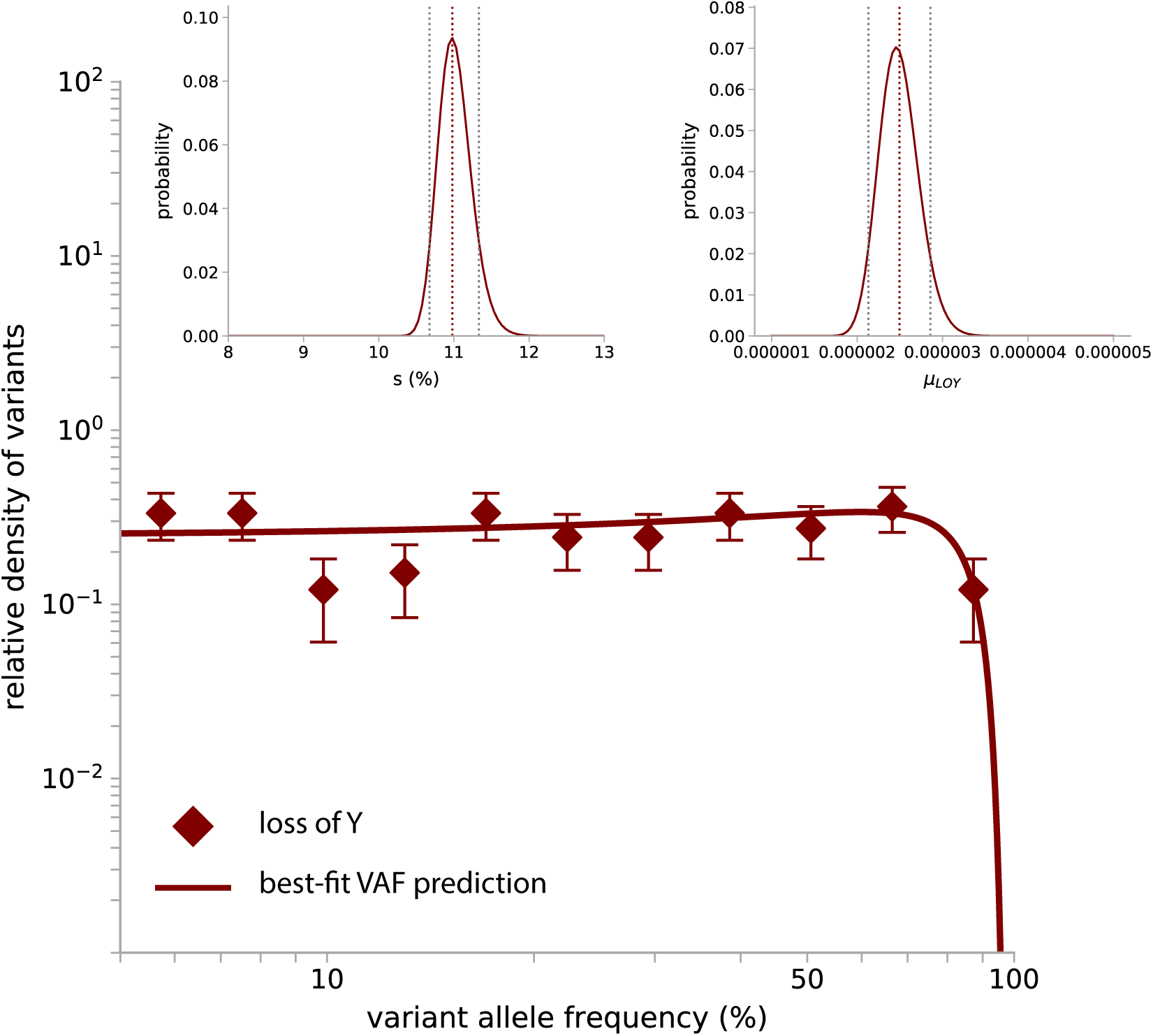
The best-fit to account for Y-loss mosaicism ^29^ predicts that loss of Y chromosome occurs at *µ*_LOY_ = 2.5 × 10^−6^ (95% CI = 5% 2.1 − 2.9 × 10^−6^) per cell per year and with fitness *s* = 11.0% per year (95% CI = 10.7% − 11.3%) (for *Nτ* = 1 × 10^5^ years). Theories are modified to account for non-diploidy and data is trimmed below VAF. Inset: Likelihood plots of the two-parameter optimization.

